# PTPN1/2 inhibits alveolar macrophage-mediated control of lung metastasis

**DOI:** 10.64898/2026.02.25.707995

**Authors:** Yue Liu, Im-Meng Sun, Marc Creixell, Jordan Brown, Samir Kharbanda, James J. Lee, Varahram Shahryari, Kayley Hake, Jacqueline O’Hara, Kenneth J. Finn, Nianxin Yang, Lolita Penland, Jiaxi Wang, Ka Man Li, John Balibalos, Aaron W. Stebbins, Patrick M. Godfrey, Po-Han Tai, Evangelia Malahias, Wenjun Kong, Nicole Fong, David Hendrickson, Shagun Gupta, Leanne Chan, Fiona McAllister, Chirag H. Patel, Marcia N. Paddock, Tuan Andrew Nguyen, Fiona A. Harding, Jonathan D. Powell

## Abstract

Metastasis remains the leading cause of cancer mortality, yet effective therapies are limited. While therapeutic responses are influenced by organ-specific immune microenvironments, strategies to pharmacologically modulate these niches remain poorly defined. Here, using the clinical-stage inhibitor ABBV-CLS-484 (AC484) as a chemical probe, we demonstrate that systemic PTPN1/2 inhibition remodels the pulmonary myeloid landscape, specifically activating alveolar macrophages (AMs) toward a tumoricidal state. Integrated single-cell/spatial transcriptomics and functional assays reveal that AC484 promotes AM accumulation within metastatic lesions, elevates their IFNγ production and responsiveness, and enhances their tumor-killing activity. Depletion of AMs diminishes the anti-metastatic efficacy of AC484. Mechanistically, inhibiting PTPN1/2 by AC484 amplifies IFNγ-STAT1 signaling in AMs, while disrupting this pathway impairs their tumor control capability. These findings delineate a distinct innate immune axis where PTPN1/2 acts as a molecular “brake” on AM activation, suggesting that pharmacologically unleashing tissue-resident macrophages offers a therapeutic strategy to overcome metastatic progression, particularly in microenvironments where adaptive immunity is insufficient.

**Significance:** AMs are potent anti-metastatic immune effectors functionally constrained by PTPN1/2. Pharmacologically inhibiting PTPN1/2 amplifies their IFNγ-STAT1 signaling and tumoricidal activity, establishing a strategy to reactivate tissue-resident immunity against metastasis.

## Introduction

Metastasis, the spread of cancer cells from a primary tumor to distant organs, is the leading cause of solid tumor-related mortality and remains a major obstacle to effective cancer therapies (1). Since the classic “seed and soil” hypothesis was proposed in 1889 (2), early anti-metastatic strategies have largely focused on tumor-intrinsic properties or on the supportive stromal elements (3). In recent decades, growing evidence has highlighted the immune system as a critical determinant throughout the entire metastatic cascade (4,5). While advances in cancer immunotherapy have demonstrated the potential to harness immunity to combat tumor spread (6), many tumors remain resistant to current therapies (7), and therapeutic efficacy often varies by metastatic site, likely reflecting distinct organ-specific immune compositions and functions (8,9). For example, the lung, a frequent site of metastasis, poses unique immunological challenges: it must constantly defend against airborne pathogens while maintaining immune tolerance to prevent excessive inflammation (10,11). This “tolerogenic” niche often suppresses anti-tumor immunity, limiting the effectiveness of T cell-centric approaches (12). As such, strategies that bypass adaptive immune resistance by re-engaging organ-specific innate immune responses represent a promising yet underexplored avenue for therapeutic intervention.

Unlike recruited myeloid cells, which are often co-opted by tumors to suppress immunity, tissue resident macrophages (TRMs) occupy specialized anatomical niches and can possess the intrinsic capacity to defend against pathogens and restrict tumor outgrowth (13,14). For example, Kupffer cells (liver resident macrophages) have been shown to limit metastatic colonization (15–17). This concept is reinforced by studies showing the anti-metastatic functions of microglia in the brain as a part of natural immune defense mechanisms (18). Therapeutic activation of microglia has been shown to enhance their tumoricidal capacity and suppress glioma growth (19). In the lung, alveolar macrophages (AMs) are the predominant resident population (10). Although AM can possess an intrinsic ability to limit lung metastasis progression (20), their therapeutic potential remains underexplored. Unlocking the tumoricidal capacity of AMs requires identifying and modulating the molecular pathways that functionally constrain their anti-tumor functions.

The non-receptor protein tyrosine phosphatases PTPN1 and PTPN2 function as critical negative regulators of cytokine signaling and intracellular immune checkpoints in both tumor and immune cells (21). By dephosphorylating key signaling molecules like JAK, PTPN1/2 limit the amplitude and duration of anti-tumor immune signaling. We and others have previously demonstrated that inhibiting PTPN1/2 promotes anti-tumor immunity and sensitizes tumors to immunotherapy primarily by enhancing CD8^+^ T cell and NK cell cytotoxicity (22–27). However, whether these phosphatases also restrict the function of tissue-resident innate immune cells in the metastatic niche remains unknown. The development of ABBV-CLS-484 (AC484), a highly selective, orally bioavailable inhibitor of PTPN1/2 (28), provides a potent chemical probe to functionally dissect this biology *in vivo*.

In this study, we establish AMs as the key effectors of lung metastatic control following PTPN1/2 inhibition. By integrating single-cell RNA sequencing (scRNA-seq) and spatial transcriptomics, we demonstrate that AC484 reshapes the pulmonary myeloid landscape during metastasis. Mechanistically, we map this response to amplified IFNγ-STAT1 signaling within AMs, which enables their potent, contact-dependent tumoricidal activity. We confirm the critical role of AMs *in vivo*, as their depletion drastically diminishes the ability of AC484 to suppress lung metastasis. Moreover, genetic or pharmacologic disruption of the IFNγ-STAT1 axis blunts metastatic suppression, identifying this pathway as a central mechanistic requirement. Collectively, our findings highlight a distinct lung resident innate immune axis, suggesting that pharmacologically unleashing AMs offers a therapeutic strategy to overcome metastatic progression.

## Results

### Inhibiting PTPN1/2 by AC484 suppresses lung metastases in spontaneous metastasis models through an immune-dependent mechanism

To study metastasis, we utilized two syngeneic mouse models with distinct genetic backgrounds: CMT167 (lung carcinoma, C57BL/6) (29) and 4T1 (breast carcinoma, BALB/c). Both models successfully developed spontaneous lung metastases, with 4T1 tumors displaying more aggressive dissemination and a higher metastatic burden (**Supplementary Figure 1A-D**). During metastatic progression, both models showed dynamic shifts in the lung immune populations, including an expansion of bone marrow-derived myeloid cells (**Supplementary Figure 1E-H**), consistent with prior reports of pre-metastatic niche formation (30,31). Notably, these immunological changes occurred early, preceding the detection of microscopic metastases.

Using the CMT167 and 4T1 models, we evaluated the effects of PTPN1/2 inhibition by AC484 on lung metastases. Treatment started on Day 10 for CMT167 and Day 7 for 4T1 post-inoculation (**Figure 1A and B**). While AC484 induced an elevation of circulating cytokines and chemokines, it did not affect primary tumor growth at the doses used (**Supplementary Figure 2A-D**). In contrast, AC484 markedly reduced parenchymal lung metastases in both models (**Figure 1C-D, F-G**).

**Figure 1.**
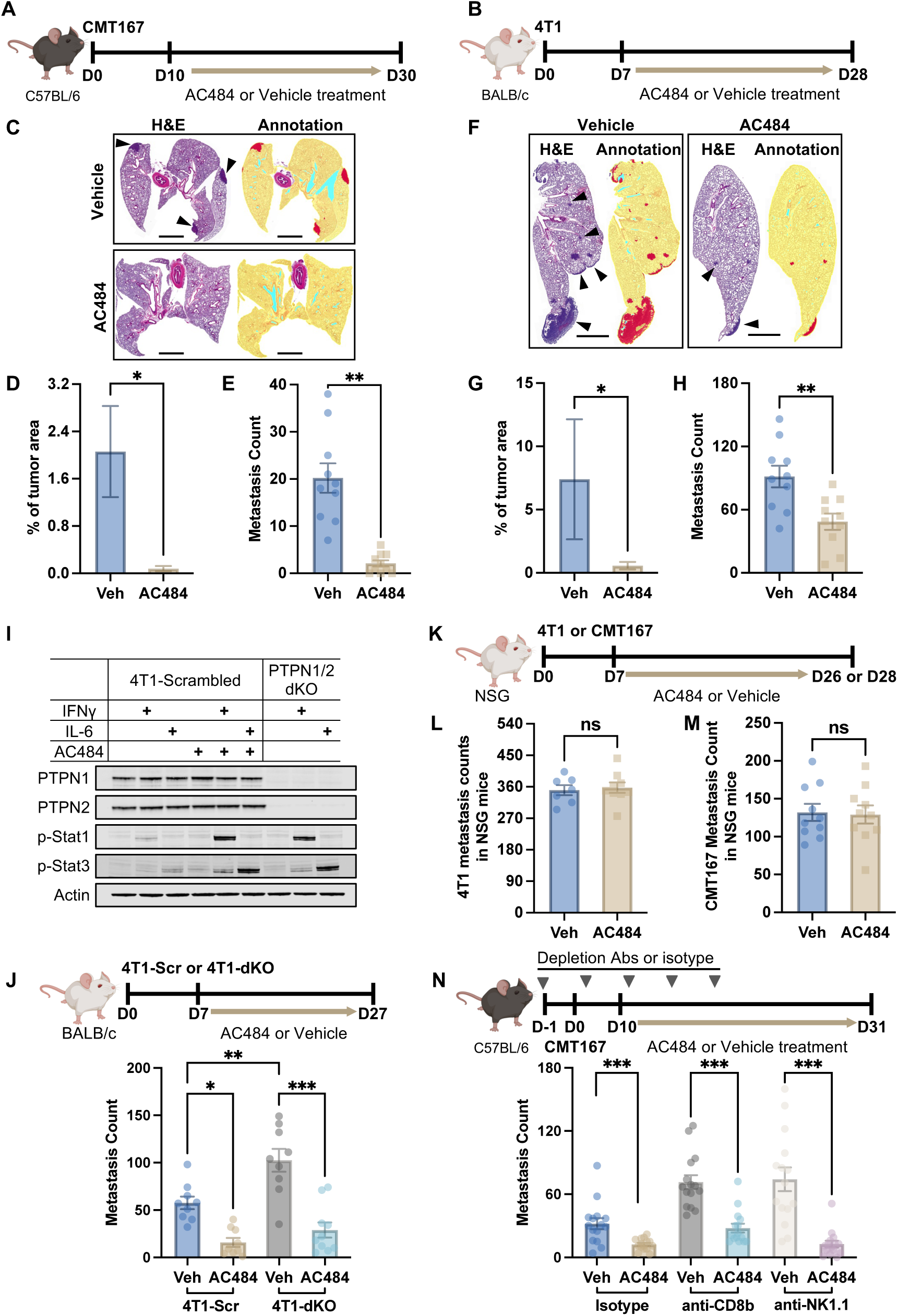
AC484 effectively suppresses lung metastasis via an immune-dependent mechanism. **A.** C57BL/6 mice were subcutaneously implanted with 1 × 10⁶ CMT167 cells and treated daily with 20 mg/kg AC484 or vehicle. **B.** BALB/c mice were implanted with 1 × 10⁵ 4T1 cells in the mammary fat pad and treated as indicated. **C.** Representative H&E-stained and analysis markup images of lungs collected at the endpoint from A. Yellow, lung parenchyma; red, metastatic tumors; light blue, glass background. Scale bar = 2 mm. **D.** Tumor area quantification from C. Shown are mean ± SEM from 3 independent sections. **E.** Visible lung surface metastasis counts in CMT167-bearing mice at the endpoint. **F.** Representative H&E-stained and analysis markup images of lungs collected at the endpoint from B. Yellow, lung parenchyma; red, metastatic tumors; light blue, glass background. Scale bar = 2 mm. Arrows point to metastatic tumor regions. **G.** Tumor area quantification from F. Mean ± SEM from 3 sections. **H.** Surface metastases count in 4T1-bearing mice at the endpoint. **I.** Western blot of PTPN1/2 double knockout (dKO) and scrambled (Scr) control 4T1 cells treated overnight with IFNγ or IL-6 (100 ng/mL), in the presence or absence of 1 μM AC484. **J.** BALB/c mice implanted with dKO or Scr 4T1 cells and treated as indicated. Lung surface metastases quantified after India ink staining. **K-M.** NSG mice implanted with 4T1 or CMT167 cells and treated as indicated. Lung metastases were visualized and quantified separately for 4T1 (**L**) and CMT167 (**M**). **N.** C57BL/6 mice were treated with depleting antibodies (anti-CD8β, anti-NK1.1, or isotype control) starting on Day -1, followed by CMT167 cell implantation and AC484 treatment. Surface metastases were quantified at the endpoint. For all bar graphs, shown are individual animals (dots), with mean ± SEM (bars). Unpaired t-test for two-group comparisons; one-way ANOVA for multiple groups: ns, *p* > 0.05; * *p* ≤ 0.05; ** *p* ≤ 0.01; *** *p* ≤ 0.001.

Assessment of lung surface tumor nodules confirmed this potent efficacy: AC484 reduced metastatic foci by over 50% compared to vehicle controls in both models (**Figure 1E-H**). To determine whether AC484 remains effective at later disease stages, we delayed treatment initiation until Day 15 post-inoculation, when a metastasis-promoting immune environment had already developed in the lungs (**Supplementary Figure 1G-H**). Remarkably, delayed AC484 administration still suppressed 4T1 lung metastases, with efficacy comparable to the early treatment cohort (**Supplementary Figure 2E**). Collectively, these findings demonstrate the robust potency of the PTPN1/2 inhibitor AC484 in controlling lung metastasis.

To parse the contribution of tumor-intrinsic versus host-mediated PTPN1/2 inhibition, we generated *Ptpn1* and *Ptpn2* double knockout (4T1-dKO) cells, along with a scrambled control (4T1-Scr) (**Figure 1I**). As PTPN1/2 function as negative regulators of JAK/STAT signaling (32), 4T1-dKO cells displayed increased phosphorylation of STAT1 and STAT3 upon IFNγ and IL-6 stimulation, respectively (**Figure 1I**). These patterns mirrored the effects of AC484 treatment in 4T1-Scr cells (**Figure 1I**), indicating that genetic loss of PTPN1/2 mimics AC484-induced changes in tumor-intrinsic signaling. 4T1-dKO and 4T1-Scr cells displayed comparable growth *in vitro* and *in vivo* (**Supplementary Figure 2F**). Compared to Scr controls, 4T1-dKO tumor cells did not exhibit reduced lung metastasis. However, systemic treatment with AC484 effectively suppressed lung metastasis in mice bearing 4T1-dKO tumors, to a similar degree as that observed in mice with 4T1-Scr (**Figure 1J**). These results indicate that the anti-metastatic activity of AC484 is not driven by tumor-intrinsic PTPN1/2 inhibition.

Given that the ability of AC484 to suppress metastasis appeared to be tumor-extrinsic, we next tested whether it required a functional immune system. We inoculated 4T1 or CMT167 tumor cells into NSG mice, an immunodeficient strain that lacks T, B, NK, and functional myeloid cells (33–36), followed by AC484 or vehicle treatment (**Figure 1K**). AC484 did not affect primary tumor growth (**Supplementary Figure 2H-J**). Most importantly, in contrast to immune-competent mice, AC484 failed to suppress lung metastasis in either model within this immunodeficient context (**Figure 1L-M**). This confirms that the anti-metastatic mechanism of PTPN1/2 inhibition is immune-dependent.

Since NK and CD8^+^ T cells are generally considered key anti-metastatic immune effectors (4), and AC484 can promote their function (28), we assessed the necessity of these cytotoxic populations for metastatic control. We depleted CD8^+^ T cells (anti-CD8β) or NK cells (anti-NK1.1) in CMT167 tumor-bearing mice. Surprisingly, depletion of neither population failed to abrogate the anti-metastatic efficacy of AC484 (**Figure 1N**), and primary tumor growth remained unaffected (**Supplementary Figure 2K**). This finding prompted us to investigate alternative immune cell types that mediate the anti-metastatic effects of AC484.

### scRNA-seq identifies anti-tumor shifts in multiple innate immune cell populations driven by AC484

To characterize the immune cells that were involved in suppressing metastasis after AC484 treatment, we performed scRNA-seq on FACS-sorted viable cells from the lungs of 4T1 tumor-bearing mice. Lungs were collected from vehicle- and AC484-treated mice (N=3 per group) at Day 14 post-tumor implantation (**Figure 2A**), a time point showing changes in the lung immune landscape but no substantial metastasis (**Supplementary Figure 1B, D, and G-H**). We generated a dataset of 76,671 cells, comprising major stromal, epithelial, and immune lineages (**Figure 2B and Supplementary Figure 3A**).

**Figure 2.**
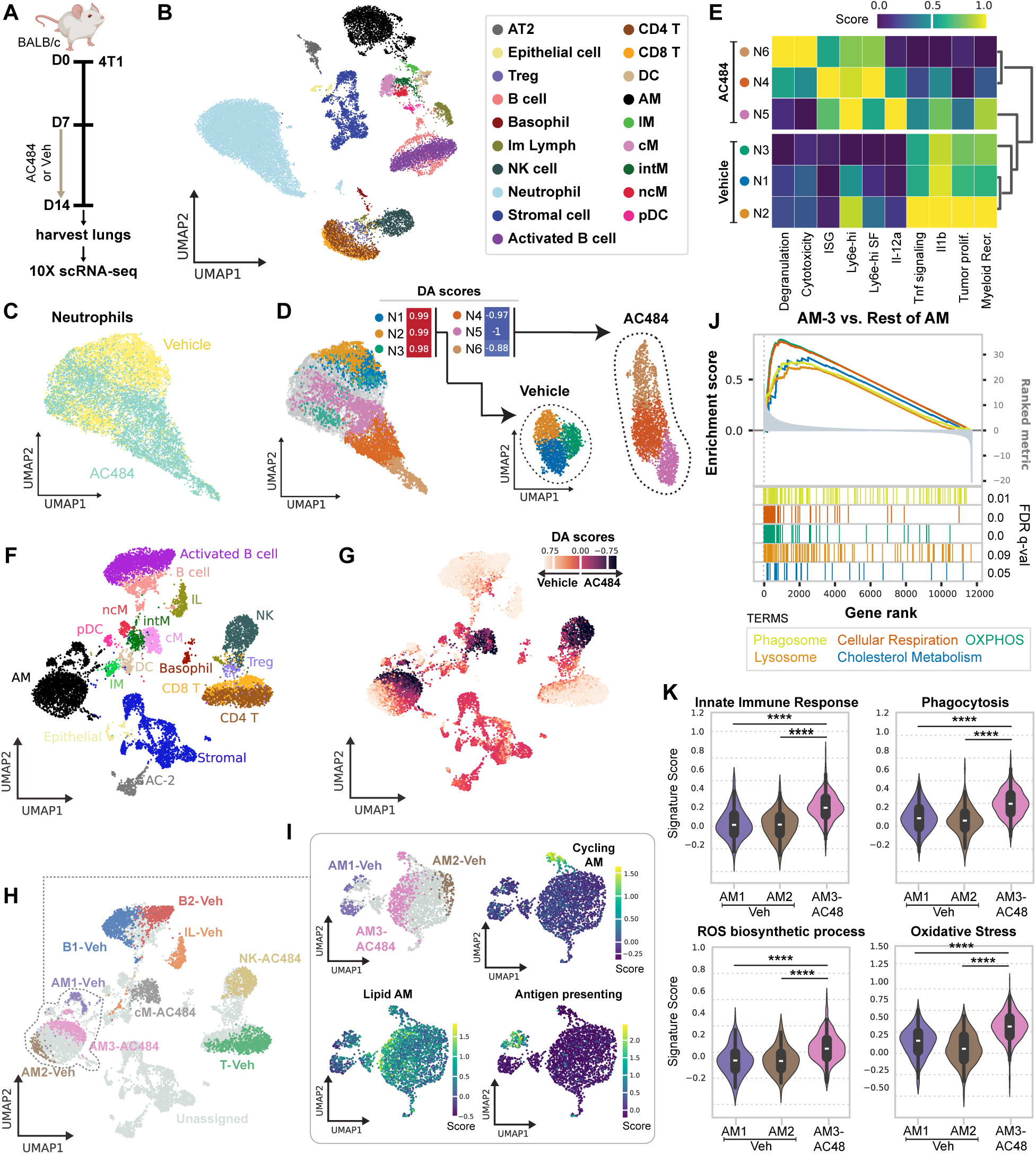
AC484 treatment results in the activation of pro-inflammatory transcriptional programs in several innate immune subpopulations during 4T1 lung metastasis. **A.** Experimental scheme showing the scRNA-seq experiment schedule. N=3 mice per treatment group. **B.** UMAP of cells that passed QC grouped by cell type. **C**. Neutrophils annotated by treatment. **D**. Differentially abundant clusters of neutrophils in each treatment group. Values indicate DAseq enrichment scores. **E**. Heatmap showing the indicated neutrophil-specific gene signatures, including normalized expression of the genes IL12a and IL1b. Functional signatures were calculated as the gene set average expression normalized to the binned expression of reference genes. **F-H.** UMAP of all cell types excluding neutrophils overlaying DA scores (**G**) or differentially abundant subpopulations (**H**). **I.** UMAP of AM subpopulations and the expression of AM-specific signatures. **J.** GSEA plot comparing enriched pathways in the AM-3 cluster (enriched following AC484 treatment) versus the rest of AMs. **K**. Violin plots showing AM-specific gene signature scores grouped by AM subpopulation. Student’s t-test. *** p ≤ 0.001.

Neutrophils were the most abundant immune cell population (**Figure 2B**), consistent with prior reports of systemic granulocytic expansion in 4T1 tumor-bearing mice (37). UMAP analysis revealed a transcriptional shift in neutrophils following AC484 treatment (**Figure 2C**). To identify treatment-specific neutrophil subpopulations, we performed differential abundance analysis using DAseq (38), and found three different neutrophil clusters per group (**Figure 2D**). Based on neutrophil-specific gene signatures (39,40), AC484-enriched clusters (N4-N6) were characterized by anti-tumor programs such as degranulation and cytotoxicity, whereas vehicle-associated clusters (N1-N3) displayed tumor-promoting signatures (**Figure 2E**). Trajectory analysis revealed that AC484 expanded immature neutrophil populations enriched for interferon-stimulated gene signatures, while vehicle treatment favored terminally differentiated neutrophils with heightened IL-1β and TNF inflammatory signaling (**Figures 2E and Supplementary Figure 3B-E**).

To focus on additional immune cell compartments, we applied DAseq excluding neutrophils (**Figure 2F-G and Supplementary Figure 3F**). This analysis revealed a significant enrichment of a subset of alveolar macrophages (AM3-AC484), along with subsets of NK cells and classical monocytes (cM) following AC484 treatment (**Figure 2G-H**).

We first focused on AM3-AC484, a subset of alveolar macrophages (AMs) with a distinct transcriptional profile enriched following AC484 treatment (**Figure 2I**). Gene set enrichment analysis (GSEA) of differentially expressed genes in AM3-AC484 revealed the upregulation of pathways related to the phagosome, lysosome, cellular respiration, oxidative phosphorylation (OXPHOS), and cholesterol metabolism (**Figure 2J**). Further analysis using AM-specific functional gene signatures (41) showed that AM3-AC484 had increased expression of signatures linked to the innate immune response, phagocytosis, and reactive oxygen species (ROS) production when compared to AM1 and AM2 (enriched in vehicle-treated mice) (**Figure 2K**). Collectively, these findings suggest a functional activation toward an anti-tumor phenotype in the AM3-AC484 cluster.

Concurrent with AM remodeling, we observed activation in other innate compartments following AC484 treatment. The NK cells enriched in AC484-treated animals (NK-AC484) were prominent sources of *Ifng*, *Gzmb*, and *Prf1*, displaying upregulated gene sets related to cytotoxicity, migration, and IFNγ responses compared to the rest of NK cells (**Supplementary Figure 4A-B**). Similarly, compared to other cMs, cM-AC484 showed increased expression of IFNγ-mediated and pro-inflammatory genes, along with metabolism and antigen presentation programs (**Supplementary Figure 4C-F**).

Collectively, our scRNA-seq analyses uncover a broad, AC484-mediated reprogramming of the lung innate immune landscape. The specific emergence of activated tissue-resident macrophages (AMs) and peripheral myeloid cells points to a coordinated response, prompting us to apply complementary approaches to determine which populations are essential for AC484-mediated metastasis control.

### Spatial transcriptomics analysis of lung sections reveals AM contribution during metastasis progression

To incorporate spatial context and pinpoint immune populations relevant to metastatic progression, we performed spatial transcriptomics on lungs harvested on Day 22 post 4T1 implantation (**Figure 3A**). At this stage, the vehicle-treated lung displayed microscopic metastatic nodules by H&E staining, whereas the AC484-treated lung did not (**Figure 3B**). Spatial cell-type mapping identified disseminated tumor cells under both conditions, but overt tumor nodules were only found in the vehicle lung section (**Figure 3C and Supplementary Figure 5A**). Consistent with our scRNA-seq data, spatial profiling of the immune landscape confirmed the prominence of myeloid cells, such as macrophages and neutrophils, in the lungs (**Supplementary Figure 5A**).

**Figure 3.**
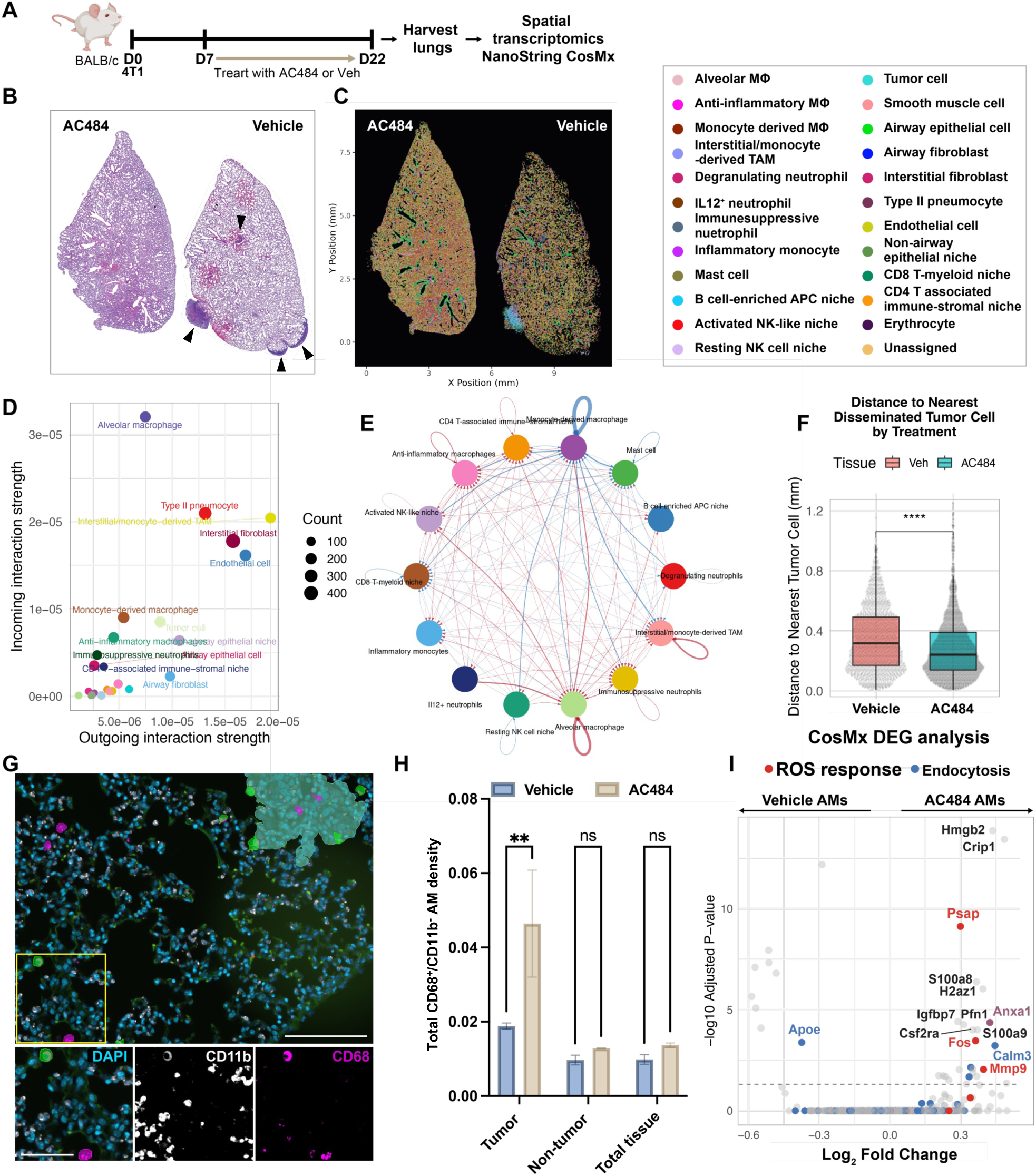
Spatial transcriptomics analysis of lung sections reveals AM contribution during metastasis progression. **A.** Experimental scheme showing BALB/c mice were implanted with 4T1 tumor cells and treated daily with AC484 or vehicle starting on Day 7 post-implantation. Lungs were harvested on Day 22 for spatial transcriptomics. **B**. H&E-stained lung sections on Day 22, corresponding to A. **C**. Spatial mapping of cell types in the serial lung section in B. **D**. CellChat analysis integrates all signaling pathways to quantify the relative communication strength among cell types. **E**. Chord diagram illustrating cell-cell communication, with outer nodules communicating to and from immune cell types. String color indicates whether the relative signal strength is increased (Red) or decreased (Blue) in the AC484-treated lung. Edge weight corresponds to the relative increase or decrease in signal strength. Arrow directionality corresponds to whether the differentially represented signaling strength is incoming or outgoing. **F**. Quantification of distances between alveolar macrophages (AMs) and disseminated tumor cells across the tissue section from C. **G.** Representative immunofluorescence images show the classification of macrophages around metastatic lesions in the lung sections of 4T1-bearing mice. Metastatic regions are semi-transparently masked in cyan; CD11b⁻ AMs are shown in magenta, and CD11b⁺ macrophages in green. Scale bar = 100 μm (upper panel) and 50 μm (lower panel). **H**. Quantification of AM density across three independent lung sections within the indicated regions. Shown are mean ± SEM. Unpaired t-test: ns, *p* > 0.05; * *p* ≤ 0.05; ** *p* ≤ 0.01; *** *p* ≤ 0.001. **I**. Differential gene expression analysis comparing AMs from AC484- and vehicle-treated lung from C. Genes are color-coded by the Gene Ontology (GO) pathway. The dotted line indicates the significance cutoff (BH adjusted p < 0.05).

Incorporating spatial constraints to quantify cell-to-cell communication events, CellChat analysis highlighted AMs as the predominant signal receivers within the metastatic lung microenvironment (**Figure 3D and Supplementary Figure 5B**). Additionally, the strength of these AM-centric interactions was significantly amplified following AC484 treatment (**Figure 3E**), prioritizing AMs as the putative drivers of the observed therapeutic response. We next quantified AM spatial proximity relative to tumor cells. AC484 treatment significantly decreased the mean distance between AMs and disseminated tumor cells (**Figure 3F and Supplementary Figure 6A**).

Hypothesizing that this proximity reflects active AM accumulation within micro-metastatic nodules, we analyzed AM distribution using CD68 immunohistochemistry (IHC) and multiplex immunofluorescence (IF). These analyses showed that AC484 selectively increased CD11b^-^ AMs, but not CD11b^+^ macrophages, specifically within metastatic tumor regions rather than non-tumor parenchyma (**Figure 3G-H and Supplementary Figure 6B-F**). Furthermore, Differential Gene Expression (DGE) analysis of AMs identified in the spatial transcriptomics demonstrated that AC484 treatment upregulated genes involved in reactive oxygen species (ROS) and endocytosis (**Figure 3I**).

To determine if tumor proximity drives phenotype, we stratified AMs into tumor-proximal and tumor-distal subsets based on the overall median AM-to-tumor distance (∼300 μm to the nearest disseminated tumor cell). In vehicle-treated lungs, the transcriptional profiles of proximal and distal AMs were similar. In contrast, AC484-treated lungs showed modest gene upregulation in proximal AMs relative to distal ones, although these changes were less pronounced than the overall transcriptional changes induced by the drug itself (**Supplementary Figure 6G-I**). These findings suggest that AC484 treatment and tumor proximity exert a combinatorial influence on the AM transcriptional state. Collectively, our sequencing data implicate AMs as central effectors in the metastatic niche and indicate that PTPN1/2 inhibition with AC484 enhances AMs’ activity.

### Alveolar macrophages display increased IFNγ production and responsiveness following AC484 treatment

As AMs are the predominant immune population of the alveolar airspace in the lung (10), we investigated how AC484 influences the local milieu by analyzing cytokines and chemokines in bronchoalveolar lavage (BAL) fluid from 4T1 tumor-bearing mice using a 13-plex analysis panel. Serum cytokine/chemokine levels were measured in parallel as a systemic reference. The most strongly upregulated cytokine in BAL following 16 days of AC484 treatment was IFNγ, followed by IFNγ-induced chemokines, such as CXCL9 and CXCL10 (**Figure 4A and B**). This effect was localized to the lung microenvironment, as no IFNγ elevation was detected in serum (**Figure 4A and C**). Notably, this local IFNγ elevation by AC484 treatment persisted through the late stages of metastasis progression (Day 29) (**Supplementary Figure 7A-C**).

**Figure 4.**
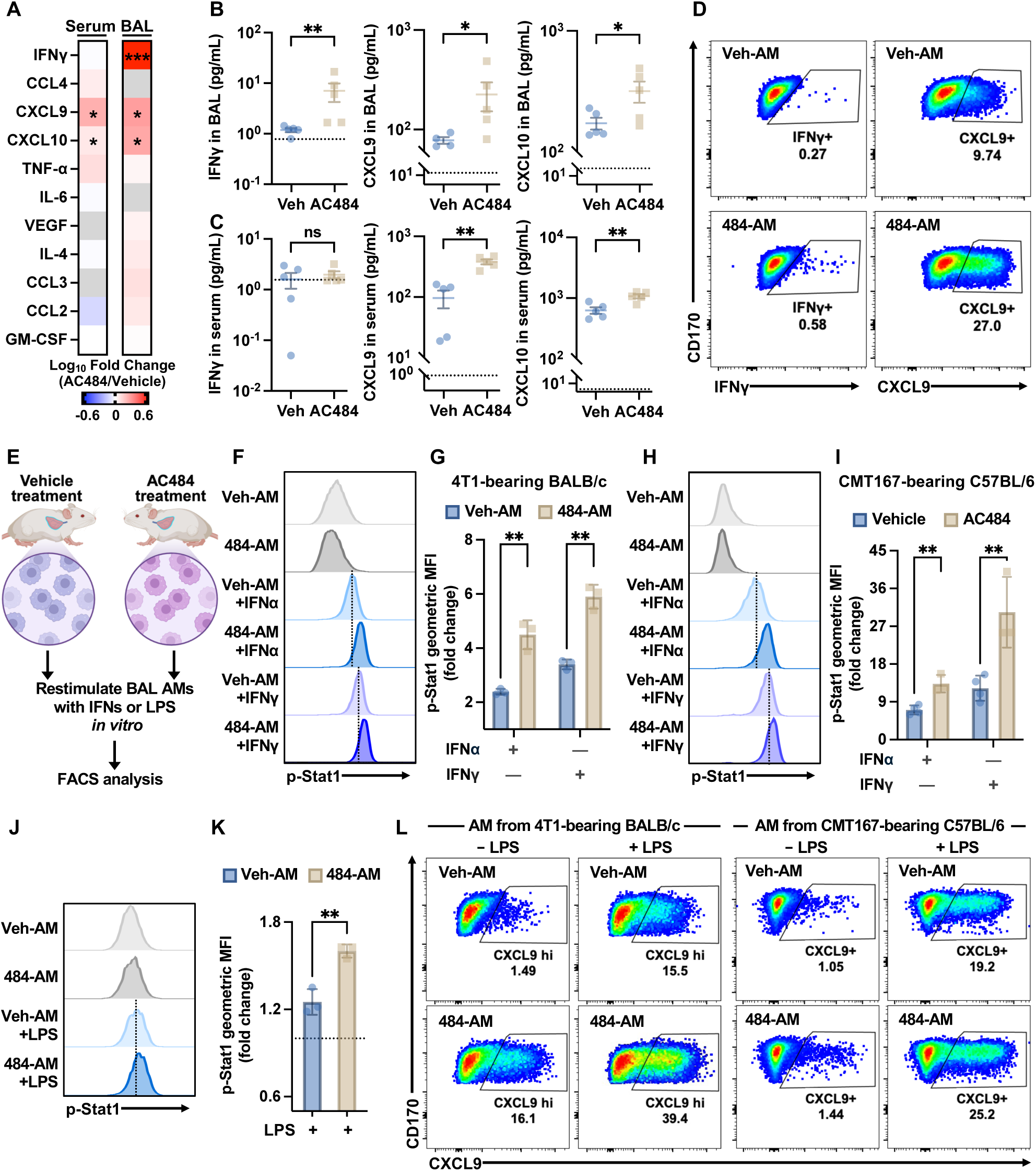
AMs display increased IFNγ production and responsiveness following AC484 treatment. **A.** Heatmap comparison of serum and BAL cytokine levels in 4T1-bearing BALB/c mice treated with AC484 or vehicle (harvested on Day 23; treatment started Day 7). Grey indicates values below the detection threshold; cytokines undetectable in both compartments were excluded. **B-C.** Quantification of specific cytokines in BAL fluid (**B**) and serum (**C**) from the dataset in A, mean ± SEM (bars). Dotted lines indicate detection limits. **D.** Intracellular IFNγ and CXCL9 staining in CD170⁺/CD11c⁺ alveolar macrophages (AMs) from AC484 (484-AM) or vehicle treated (Veh-AM) 4T1-bearing BALB/c mice. **E.** Experimental scheme showing AMs isolated from AC484 or vehicle-treated tumor-bearing mice were stimulated *ex vivo* with interferons or LPS. **F-I.** The level of p-STAT1 (Y701) in AMs following stimulation with IFNα or IFNγ (150 ng/mL, 15 minutes). **F, H.** Representative histograms of AMs from 4T1-bearing (**F**) and CMT167-bearing (**H**) mice. **G, I.** Fold change of p-STAT1 geometric MFI normalized to unstimulated controls, corresponding to F and H. **J-K.** The level of p-STAT1 (Y701) in AMs from 4T1-bearing animals with or without LPS stimulation (2.5 µg/mL, 2.5 h). Shown are representative histograms (**J**) and a normalized geometric MFI fold change (**K**). **L.** Intracellular CXCL9 levels in AMs collected from AC484 or vehicle treated tumor-bearing mice, with or without *ex vivo* LPS stimulation. For all bar graphs, data represent individual animals (dots) and mean ± SEM (bars). Statistical significance was determined using unpaired t-test: ns, *p* > 0.05; * *p* ≤ 0.05; ** *p* ≤ 0.01; *** *p* ≤ 0.001.

To investigate whether AMs are a source of IFNγ, we harvested BAL AMs from 4T1 tumor-bearing mice and examined their cytokine expression. Our results demonstrated that AMs from AC484-treated mice showed increased levels of IFNγ and CXCL9 compared to those from vehicle-treated mice (**Figure 4D**), which may at least partially account for the elevated IFNγ detected in BAL. Since PTPN1 and PTPN2 can act as negative regulators of interferon (IFN)-JAK/STAT signaling (42), their inhibition is expected to amplify this pathway. To test this hypothesis in AMs, we stimulated them *in vitro* with type I (IFNα) or type II (IFNγ) interferons (**Figure 4E**). AMs from AC484-treated mice (484-AMs) displayed significantly higher levels of p-STAT1 (Y701) than those from vehicle-treated controls (Veh-AMs) (**Figure 4F-I**). This confirms successful *in vivo* target engagement of AC484 and demonstrates enhanced IFN responsiveness in AMs upon PTPN1/2 inhibition. Together, these results suggest that in AC484-treated mice, AMs have heightened sensitivity when stimulated by IFNγ produced either by AMs or other immune cells in the local lung microenvironment (e.g., BAL).

To further validate this IFNγ primed state, we stimulated AMs with lipopolysaccharide (LPS). While LPS alone typically does not induce STAT1 phosphorylation, it triggers a robust STAT1 response in macrophages pre-exposed to IFNγ (43). Consistent with this paradigm, Veh-AMs responded only modestly to LPS, whereas 484-AMs exhibited robust STAT1 (Y701) phosphorylation and markedly higher production of CXCL9 (**Figure 4J-L**), a STAT1-driven chemokine (44). Collectively, these data indicate that PTPN1/2 inhibition by AC484 in tumor-bearing mice enforces an IFNγ-primed state in AMs, enabling amplified STAT1 activation and downstream signaling.

### AMs are essential mediators of AC484-induced metastasis control

To directly link AM activation to tumor control, we cocultured BAL AMs from tumor-bearing mice with matched tumor cell lines (4T1 or CMT167) in the presence or absence of AC484 (**Figure 5A**). While coculturing AMs with tumor cells alone resulted in a modest increase of p-STAT1 levels in AMs, the addition of AC484 in the coculture significantly enhanced STAT1 phosphorylation (**Figure 5B-E**). Functionally, this translated to enhanced cytotoxicity: AC484 significantly increased AM-mediated tumor cell killing in both models at 72 hours, with a stronger effect observed in 4T1 compared to CMT167 (**Figure 5F-G**).

**Figure 5.**
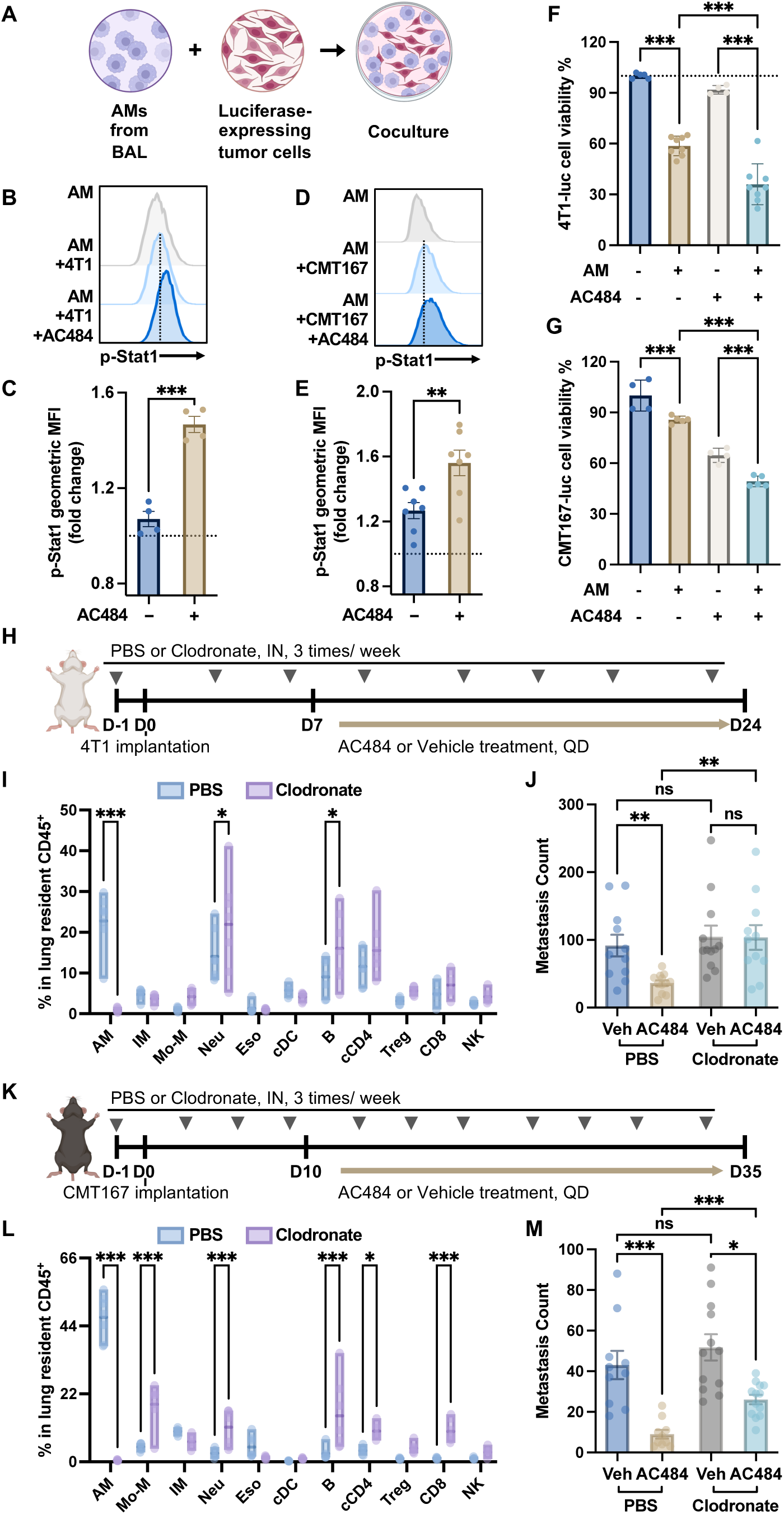
AMs are critical for the ability of AC484 to inhibit metastasis. **A.** Experimental scheme showing alveolar macrophages (AMs) isolated from tumor-bearing mice were cocultured with tumor cells *in vitro*. **B-E.** The level of p-STAT1 (Y701) in AMs cocultured with tumor cells ± AC484 (100 nM). Representative histograms of CD170⁺/CD11c⁺ AMs (**B, D**) and geometric MFI fold change normalized to non-cocultured controls (**C, E**) are shown for cells derived from 4T1-bearing (B, C) or CMT167-bearing (D, E) mice. **F-G.** Viability of 4T1 (**F**) and CMT167 (**G**) cells after 3-day coculture with or without AMs, in the presence or absence of AC484 (300 nM). **H-J.** Effects of AM depletion on metastasis in the 4T1 model. **H.** Experimental scheme showing BALB/c mice were administered intranasally with clodronate (50 µL/mouse) or PBS liposomes starting Day -1, followed by 4T1 tumor implantation and the indicated treatment. **I.** Flow cytometry analysis of lung-resident CD45⁺ immune cells the day after the 3rd clodronate dose. **J.** Lung surface metastases were quantified at the endpoint in mice from H. **K-M.** Effects of AM depletion on metastasis in the CMT167 model. **K.** Experimental scheme for intranasal clodronate or PBS liposomes (50 µL/mouse) administration and treatment schedule in CMT167-bearing C57BL/6 mice. **L.** Flow cytometry analysis of lung-resident CD45⁺ immune cells. **M.** Lung surface metastases were quantified at the endpoint in mice from K. For all bar graphs, data represent individual animals (dots) and mean ± SEM (bars). Unpaired t-test for two-group comparisons; one-way ANOVA for more than two groups: ns, *p* > 0.05; * *p* ≤ 0.05; ** *p* ≤ 0.01; *** *p* ≤ 0.001.

To investigate the role of AMs *in vivo*, we treated mice with intranasal administration (IN) of clodronate liposomes (**Figure 5H**). This method efficiently depleted AMs while sparing other lung immune populations, including interstitial macrophages (IMs), T cells, and NK cells (**Figure 5I**). The significant decrease in the AM percentage in the lung is likely compensated by an increase in other cell populations, such as neutrophils, consistent with findings from other studies (45). Local AM depletion did not affect primary tumor growth, AC484’s effect on tumor size, or overall mouse health as measured by body weight (**Supplementary Figure 8A-B**). In the vehicle-treated group, the lung metastatic burden was not impacted by clodronate alone. However, AM depletion abolished the ability of AC484 to suppress metastasis in the 4T1 model (**Figure 5J**), demonstrating that AMs are the primary cell type responsible for AC484’s anti-metastasis properties in this model.

The 4T1 model has been characterized by strong myeloid dependency, weak lymphocyte responses, and resistance to ICB (46). Consistent with these features, anti-PD1 monotherapy failed to control tumor growth or metastasis and provided no additional benefit when combined with AC484 (**Supplementary Figure 8C-E**). The lack of an anti-PD1 response in our 4T1 model aligns with the limited role of functional T cells, particularly CD8⁺ T cells, raising the possibility that the strong reliance of the 4T1 model on AMs may stem from its minimal lymphocyte contribution. Unlike 4T1, CMT167 primary tumors respond to AC484 **+** anti-PD1 combination therapy (**Supplementary Figure 8F-H**), indicating successful activation of anti-tumor immunity likely mediated by T cells. Anti-PD1 monotherapy also reduced CMT167 lung metastases (**Supplementary Figure 8H**), further supporting a prominent role for lymphocytes in this model. To determine if AMs remain critical for AC484’s efficacy against metastasis in this context, we depleted AMs in CMT167-bearing mice (**Figure 5K**). Notably, the depletion of AMs in the lungs of CMT167-bearing mice led to a greater upregulation of T cells than was observed in 4T1-bearing BALB/c mice (**Figure 5L**). Although AM depletion did not affect subcutaneous primary tumors or AC484’s effect on them (**Supplementary Figure 8I-J**), it significantly impaired AC484’s anti-metastatic capability in CMT167-bearing mice, roughly tripling lung metastases compared to non-depleted, AC484 treated animals (**Figure 5M**). Thus, AMs remain critical mediators of AC484’s anti-metastatic activity, although their relative contribution can vary across tumor models depending on their broader immune context.

To distinguish the contributions of AMs from that of recruited bone marrow-derived macrophages, we treated mice intraperitoneally (IP) with antibodies against colony-stimulating factor 1 receptor (CSF1R), a receptor essential for the survival and differentiation of macrophages derived from bone marrow precursors (47,48). Consistent with the previous reports (49,50), systemic anti-CSF1R treatment reduced bone marrow-derived IMs, while having less impact on lung-resident AMs (**Supplementary Figure 9A-B**). Anti-CSF1R alone had no noticeable impact on 4T1 primary tumor growth or metastasis (**Supplementary Figure 9C-D**). Importantly, AC484 treatment significantly reduced metastasis regardless of CSF1R blockade (**Supplementary Figure 9D**). These data reinforce lung-resident AMs as the primary mediators of AC484’s anti-metastatic effect.

### IFNγ/STAT1 signaling is critical for AC484-enhanced anti-tumor activity of AMs

Our results so far identified AMs as critical effectors of AC484-mediated metastasis control, with IFNγ signaling emerging as a key pathway, we next sought to delineate the functional contribution of IFNγ signaling to this process. We cocultured AMs isolated from 4T1- or CMT167-bearing mice with tumor cells in the presence or absence of recombinant IFNγ, AC484, or both. In the absence of AMs, 1.6 ng/mL IFNγ induced only modest direct cytotoxicity against tumor cells, an effect further augmented by AC484. However, the presence of AMs significantly drove tumor killing in an IFNγ dose-dependent manner, and AC484 treatment further amplified this AM-mediated cytotoxicity (**Figure 6A-B**). These results suggest that while IFNγ endows AMs with tumoricidal capacity, PTPN1/2 inhibition sensitizes these cells to the cytokine, enhancing their killing potential even at limiting IFNγ concentrations.

**Figure 6.**
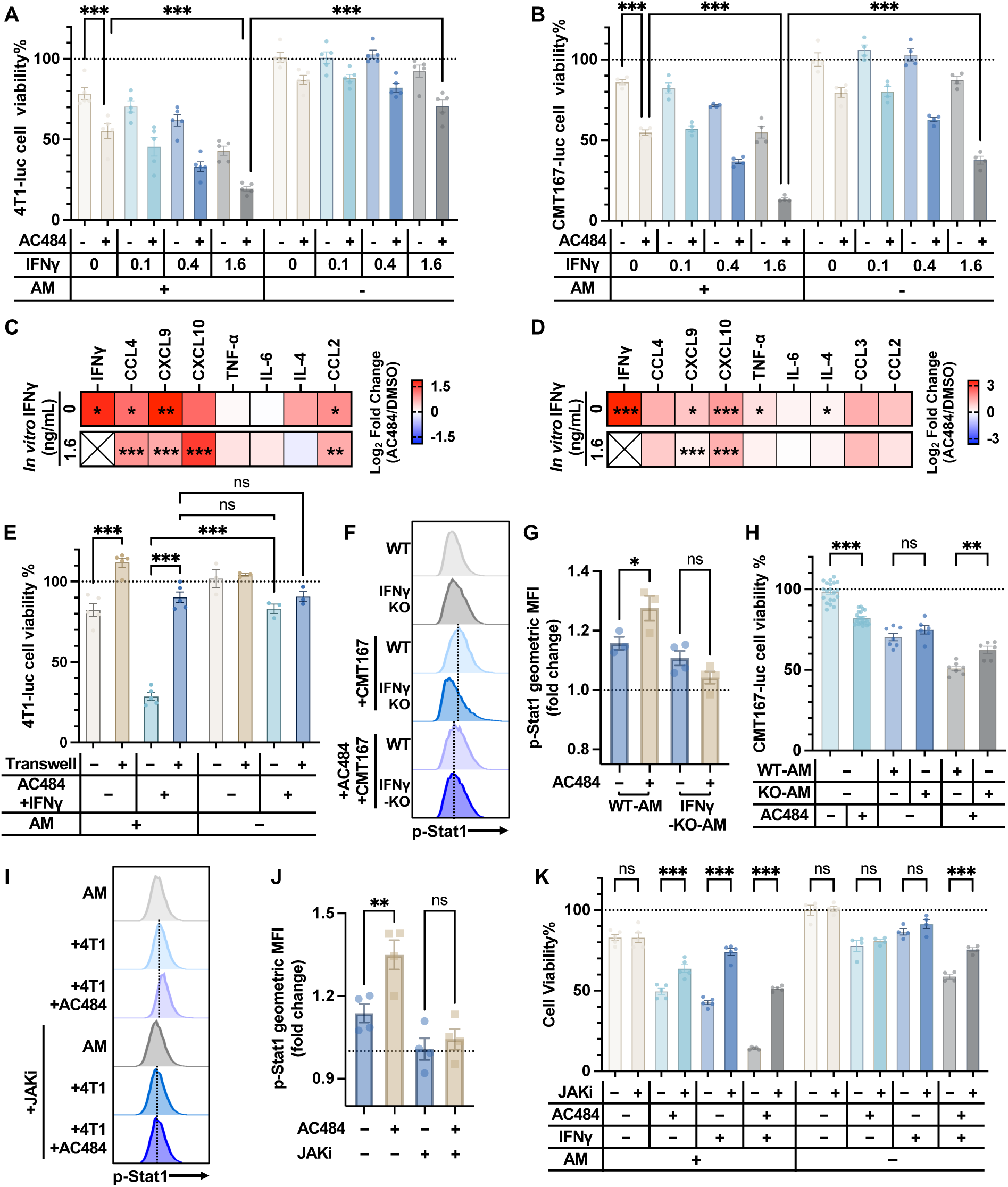
IFNγ-STAT1 signaling is involved in the anti-tumor activity of AMs enhanced by AC484. **A, B.** Viability of 4T1 (**A**) and CMT167 (**B**) cells after a 3-day coculture with or without AMs, in the presence of AC484 (50 nM) and/or IFNγ at indicated concentrations (ng/mL). **C, D.** Comparison of cytokine levels in conditioned medium collected from AMs cocultured with 4T1 (**C**) or CMT167 (**D**) tumor cells, supplemented with AC484 or DMSO controls. Cross makes indicate values beyond the detection range. Cytokines in the 13-plex panel that did not reach the detection limit in either condition were not plotted. **E**. Viability of 4T1 cells after coculture with or without AMs, comparing direct cell-to-cell contact versus separation by a transwell insert. **F**. Representative histograms of p-STAT1 (Y701) in AMs from wild-type (WT) or IFNγ knockout (KO) mice after 48 h coculture with CMT167 cells. AMs cultured alone serve as controls. **G**. Analysis of the fold change in p-STAT1 geometric MFI normalized to non-cocultured AMs from F. **H**. Viability of CMT167 cells after coculture with or without AMs harvested from WT or IFNγ KO mice, in the presence or absence of AC484. **I**. Representative histograms of p-STAT1 (Y701) in AMs after 24 h coculture with 4T1 tumor cells, with AC484 (100 nM) and/or the JAK inhibitor ruxolitinib (3 μM). AMs cultured alone with DMSO served as controls. **J**. Analysis of the fold change in p-STAT1 geometric MFI, normalized to DMSO-treated, non-cocultured control AMs, from experiments in I. **K**. Viability of 4T1 cells after coculture with or without AMs, in the presence or absence of AC484 (100 nM) and/or ruxolitinib (3 μM). For all bar graphs, data represent individual animals (dots), with mean ± SEM (bars). Unpaired t-test for two-group comparisons; one-way ANOVA for more than two groups: ns, *p* > 0.05; * *p* ≤ 0.05; ** *p* ≤ 0.01; *** *p* ≤ 0.001.

To explore the signaling basis, we analyzed cytokine and chemokine levels in the coculture supernatants. In the absence of exogenous IFNγ, AC484 treatment robustly increased the secretion of endogenous IFNγ as well as the IFNγ-responsive chemokines CXCL9 and CXCL10, confirming that PTPN1/2 inhibition promotes IFNγ production and potentially drives an autocrine activation loop in AMs (**Figure 6C-D and Supplementary Figure 10A-B**). Furthermore, even upon the addition of recombinant IFNγ, AC484 continued to boost the secretion of downstream chemokines (**Figure 6C-D and Supplementary Figure 10C-D**). Collectively, these findings suggest a dual action of AC484: it not only promotes IFNγ-production, driver of AM anti-tumor immunity, but also amplifies the subsequent IFNγ response in AMs. This two-pronged modulation effectively drives AMs toward a hyper-responsive, tumoricidal state.

To define the physical requirements of AM-mediated cytotoxicity, we separated AMs and tumor cells using a transwell system. While direct contact facilitated potent tumor killing, the anti-tumor effects of the AM + IFNγ + AC484 combination were significantly blunted when cells were physically separated (**Figure 6E**). Conversely, the modest direct cytotoxicity of the IFNγ + AC484 treatment on tumor cells remained unaffected by the transwell (**Figure 6E**). These findings support a contact-dependent mechanism for AM-mediated tumor killing.

We next asked whether IFNγ signaling is required for AC484 to enhance AM’s anti-tumor activity. To test this, we harvested BAL AMs from CMT167-bearing wild-type (WT) or IFNγ-deficient (KO) mice and cocultured them with tumor cells. While AC484 upregulated p-STAT1 (Y701) in WT-AMs during coculture, it failed to boost p-STAT1 in IFNγ-KO-AMs (**Figure 6F-G**). Functionally, while IFNγ-KO-AMs retained baseline tumor-killing capacity similar to WT-AMs, the ability of AC484 to enhance this killing was substantially diminished (**Figure 6H**). To confirm the downstream signaling node, we pretreated AMs with the JAK1/2 inhibitor ruxolitinib (JAKi). JAK inhibition blocked the AC484-induced upregulation of p-STAT1 and diminished the enhancement of AM tumor-killing activity by AC484 and/or IFNγ (**Figure 6I-K**). These *in vitro* findings establish IFNγ-JAK/STAT1 axis as the critical signaling node by which AC484 amplifies AM’s anti-tumor activity, although this does not preclude the involvement of parallel signaling mechanisms.

Finally, we validated the role of the IFNγ signaling axis *in vivo*. Blocking IFNγ via neutralizing antibodies or genetic ablation (IFNγ KO mice) had no impact on primary tumor growth or cause systemic toxicity, regardless of AC484 treatment (**Figure 7A-B, D-E, and Supplementary Figure 11A-B**). However, in the 4T1 model, IFNγ neutralization phenocopied AM depletion, abolishing the anti-metastatic efficacy of AC484 (**Figure 7C**). In IFNγ KO mice bearing CMT167 tumors, baseline lung metastases were markedly higher than in WT controls. More importantly, the efficacy of AC484 was severely compromised in in the absence of host IFNγ, achieving only a ∼20% reduction in metastasis compared to >85% in WT controls (**Figure 7F**). Collectively, these findings identify IFNγ signaling as an essential pathway for the ant-metastatic efficacy of PTPN1/2 inhibition, without excluding the potential contribution of additional pathways to the overall therapeutic effect.

**Figure 7.**
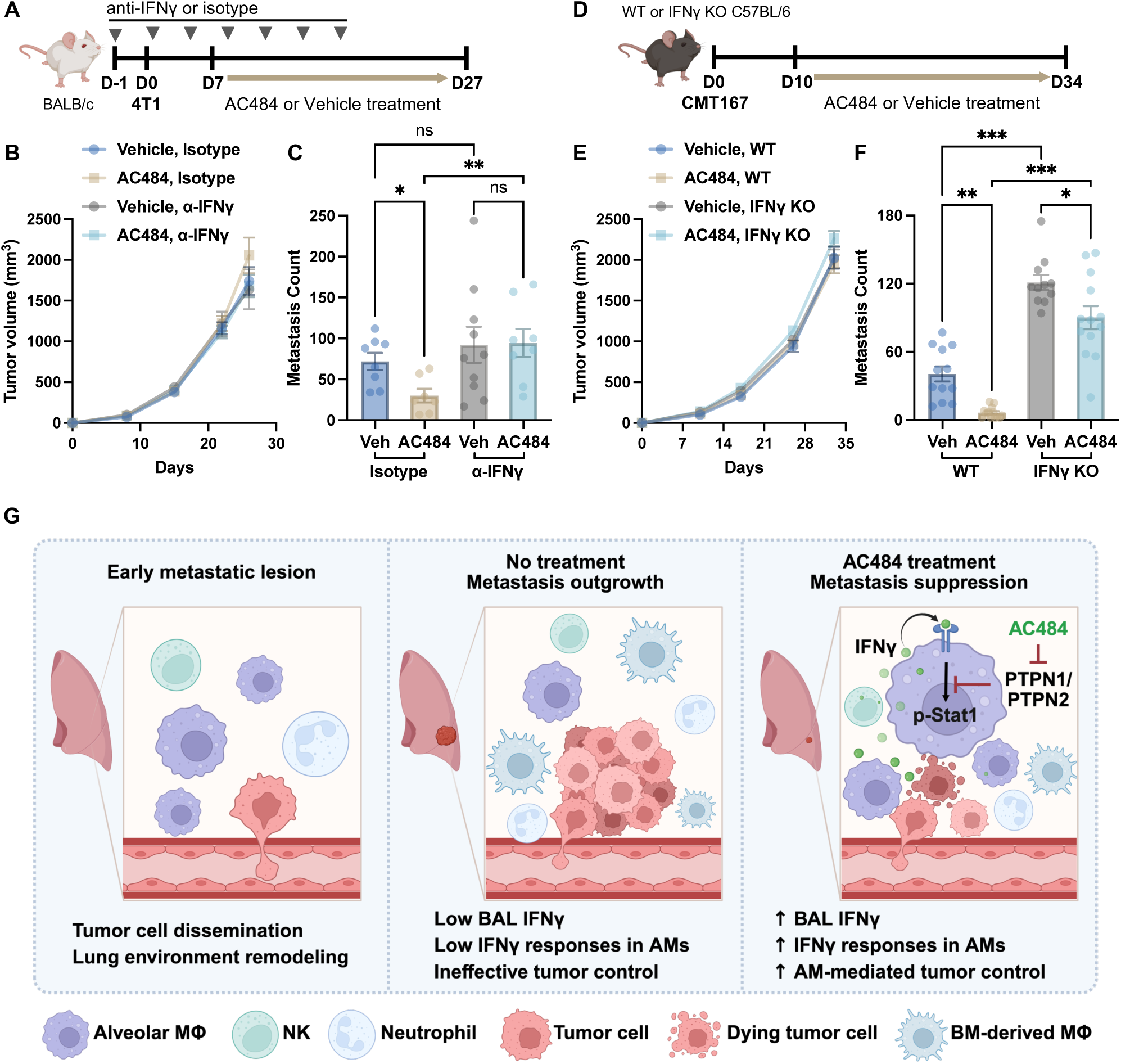
IFNγ signaling is required for the anti-metastatic activity of AC484. **A-C**, Effect of IFNγ neutralization in the 4T1 model. **A.** Experimental scheme showing BALB/c mice received anti-IFNγ antibody (200 µg/mouse) or isotype control on Day -1, followed by 4T1 implantation and the indicated treatment. **B.** Primary tumor growth kinetics are shown as mean ± SEM. **C.** Quantification of lung surface metastases at the endpoint. **D-E,** Effect of host IFNγ deletion in the CMT167 model. **D.** Experimental scheme showing IFNγ knockout (KO) or wild-type (WT) mice implanted with CMT167 cells and treated as indicated. **E.** Primary tumor growth kinetics are shown as mean ± SEM. **F.** Lung surface metastases were quantified at the endpoint in CMT167-bearing mice. **G.** Graphical summary showing that AC484 suppresses lung metastasis by increasing local IFNγ in the lung microenvironment and sensitizing AMs to IFNγ signaling, thereby enabling them to control metastatic tumor cells. Figure created with BioRender.com. For metastasis counting, shown are individual animals (dots) with mean ± SEM (bars). One-way ANOVA test: ns, *p* > 0.05; * *p* ≤ 0.05; ** *p* ≤ 0.01; *** *p* ≤ 0.001.

## Discussion

In this study, we employed two spontaneous preclinical lung metastasis models to capture distinct immune landscapes: the orthotopic 4T1 breast cancer model, characterized by aggressive metastatic progression and profound myeloid cell accumulation, and the heterotopic flank CMT167 lung cancer model, which displays slower metastasis development, fewer myeloid cells, and a greater response to ICB. We intentionally selected the C57BL/6-based CMT167 model to complement the BALB/c-based 4T1 model, which is known for its exaggerated granulocytic response (51,52). Notably, despite their different tissues of origin, genetic backgrounds, and immune microenvironments, PTPN1/2 inhibition via AC484 consistently suppressed lung metastases in both models.

Our data identify AMs as the central effectors of the anti-metastatic immunity induced by PTPN1/2 inhibition (**Figure 7G**). These findings highlight the specific importance of AMs, a tissue-resident myeloid lineage originating from yolk sac erythro-myeloid progenitors (EMPs) with self-renewal capacity (53–55). While prior studies have shown that genetic deletion of *Ptpn1* or *Ptpn2* enhances inflammatory activation in monocytes-derived macrophages (56–59), potentially supporting anti-tumor responses (28,60). Since monocytes-derived macrophage populations (including BMDMs and TAMs) primarily depend on CSF1R signaling for differentiation and survival (47–50,61), one might predict that CSF1R blockade would impact AC484’s efficacy. However, AC484 retained its anti-metastatic potency even under CSF1R blockade. These observations underscore the functional specializations of macrophage subsets and highlight the unique anti-metastasis properties of AMs driven by PTPN1/2 inhibition. As most organs harbor specialized niches of tissue-resident macrophages (TRMs) (14,62,63), our findings suggest a potential paradigm where PTPN1/2 inhibition could reprogram TRM populations, particularly in other metastatic-prone sites, such as the liver, brain, and bone.

TRMs occupy a complex duality in cancer, often described as having a paradoxical role in either promoting tumor growth or supporting anti-tumor responses (12,64–66). Here, we demonstrate that AMs, a TRM population in the lung, can be modulated by PTPN1/2 inhibition to enhance their tumoricidal potential, at least in part through increased IFNγ signaling. Prior studies have reported that stimulation with LPS and/or IFNγ commonly polarizes AMs toward an anti-tumor phenotype (67–69). IFNγ itself has long been recognized as a potent anti-tumor cytokine in the pulmonary environment. For instance, inhaled IFNγ can suppress lung metastases in mice and enhance oxygen radical production by AMs in humans (70,71). Subsequent research revealed that IFNγ produced during an influenza infection can induce transcriptional changes in AMs, enabling them to acquire tumoricidal properties (69). Building on this, we find that inhibition of the phosphatases PTPN1 and PTPN2, negative regulators of the IFNγ-JAK-STAT1 signaling axis (21,72), significantly enhances IFNγ responsiveness in AMs and promotes their tumor control capacity (**Figure 7G**). Importantly, using AC484 to enhance this pathway represents a distinct therapeutic approach compared to direct cytokine delivery or microbial stimulation. This pharmacologic amplification of IFNγ signaling provides a complementary and potentially more tunable strategy to activate tissue-resident innate immunity.

We observed elevated IFNγ in AM-tumor cocultures and in BAL fluids following AC484 treatment. While AMs are capable of producing IFNγ (73,74), a capacity further amplified by PTPN1/2 inhibition, our findings suggest that AMs serve as both an important source and target of IFNγ signaling in the AC484-mediated control of lung metastasis (**Figure 7G**). However, contributions from other immune cells cannot be excluded and are in fact likely, given the known mechanisms of AC484. Previous studies have shown that IFNγ sources in the lungs are diverse and context-dependent, ranging from phagocytes T and NK cells (69,75–77). Consistent with this, our scRNA-seq analysis revealed increased IFNγ transcripts in NK cells following AC484 treatment in the 4T1 model. Moreover, our data indicate that AC484 enhances IFNγ signaling in neutrophils and monocytes, cell types typically associated with pro-tumor activity (78,79). As IFNγ responses in these cells have also been linked to anti-tumor outcomes (40,80), AC484 may reprogram these myeloid cells to support the broader anti-metastatic potential, either directly or indirectly. Taken together, our findings imply that PTPN1/2 inhibition engages a cooperative immune network in the lung. Future studies are needed to define how AC484 shapes the immune landscape across distinct organ environments and identify additional cellular mediators and cytokine pathways that drive therapeutic efficacy.

Our findings highlight the potential of targeting intracellular “brakes”, such as PTPN1/2, to unleash context-specific immune responses, positioning this as a tractable strategy to potentiate anti-metastatic immunity. While IFNγ plays a critical role in the AC484-mediated anti-metastasis setting, PTPN1/2 can also regulate additional cytokine pathways, including IL-6, IL-2, and Type I IFNs (32). These pathways are involved not only in anti-tumor responses but also in antimicrobial defense and lung homeostasis (11,81). Therefore, AC484 may have broader immunomodulatory potential in infectious or inflammatory lung diseases. Future studies are needed to dissect the broader signaling networks governing AM plasticity and identify the most therapeutically actionable pathways across disease contexts, enabling rational combination strategies that maximize efficacy while preserving tissue integrity.

In summary, our data demonstrate that PTPN1/2 inhibition alters the pulmonary immune microenvironment to suppress metastasis, identifying AMs as central effectors. With AC484 currently in Phase 1 trials, these findings highlight a timely opportunity to complement T-cell-focused immunotherapies by awakening tissue-resident innate immunity. As an orally bioavailable small molecule, AC484 offers a tractable strategy to complement curative-intent surgeries or adjuvant therapies, potentially controlling lung metastases in high-risk patients. Finally, the ability of AC484 to potentiate cytokine signaling suggests broader utility beyond oncology, potentially enhancing infectious disease management where robust cytokine responses are required.

## Material and Methods

### Cell lines and cell culture

4T1 breast cancer cell line (CRL-2539, ATCC) and CMT167 lung cancer cell line (Clone of CMT 64, Sigma Aldrich) were respectively cultured in RPMI-1640 and Dulbecco’s Modified Eagle Medium (Gibco) supplemented with 10% Fetal Bovine Serum and 1% Penicillin-Streptomycin (complete medium).

The 4T1-Luc and CMT167-Luc cell lines were established through lentiviral transduction with vectors encoding Firefly luciferase. Single-cell clones of luciferase-transduced cells were cultured and tested using the ONE-Glo™ EX Luciferase Assay (Promega). Ten clones with high and consistent luciferase activity were selected and pooled to establish the final cell line.

*Ptpn1/2* double knockout 4T1 cells were generated using CRISPR/Cas9 ribonucleoprotein (RNP) (Synthego) following the manufacturer’s protocol. Briefly, 180 pmol of sgRNA was combined with 20 pmol of Cas9 protein in SE Cell Line Nucleofector Solution (Lonza), mixed, and incubated for 15 minutes at room temperature to form the RNP complex. 4T1 cells were mixed with the RNP complex and electroporated using the DT-130 program on a Lonza 4D-Nucleofector system. Gene knockout efficiency was validated via amplicon sequencing and Western Blot. The sequences of sgRNA and primers are listed in Supplementary Table 1. Regular *Mycoplasma* testing confirmed the absence of contamination in all cells.

### Mouse models and *in vivo* treatments

C57BL/6, BALB/c, NSG (strain #005557), and B6.129S7-*Ifngtm1Ts*/J (strain # 002287) mice were purchased from Jackson Laboratory. All mice were maintained at Calico Life Sciences and used between the ages of 6 to 18 weeks. All studies were approved by Calico’s Institutional Animal Care and Use Committee (IACUC). Humane euthanasia was performed in accordance with IACUC protocol when tumor volume exceeded 3,000 mm³, if body weight loss was 20% or more, or if the body condition score (BCS) was 2 or less.

Spontaneous lung metastasis models were established using two cell lines. For the 4T1 model, female BALB/c mice or NSG mice were implanted with 1×10^5^ cells into the mammary fat pad. For the CMT167 model, C57BL/6 mice or other indicated strains received a subcutaneous implantation of 1×10^6^ cells into the right flank. Tumor volume was calculated as 0.5×length×width×width.

The PTPN1/2 inhibitor ABBV-CLS-484 (AC484) was provided by AbbVie and formulated for oral gavage (*p.o.*) as previously described (28). Mice received AC484 daily at 20 mg/kg, with treatment initiating on Day 7 post-4T1 tumor inoculation and on Day 10 post-CMT167 tumor implantation or as otherwise indicated.

For immunotherapies, mice received intraperitoneal (*i.p.*) injections of 250 µg anti-PD-1 (CD279) (clone RMP1-14, BioXcell) every three days. For cell depletion, mice were administered *i.p*. injections of 200 µg anti-CD8β (clone 53.5.8, BioXcell) every four days to deplete CD8⁺ T cells, 200 µg anti-NK1.1 (clone PK136, BioXcell) every four days to deplete NK cells, 500 µg anti-CSF1R (clone AFS98, BioXcell) every other day to deplete interstitial macrophages (49,82), and 200 µg anti-IFNγ (clone XMG1.2, BioXcell) every three days to neutralize IFN-γ. Corresponding isotype controls, including rat IgG2a (Clone 2A3, BioXcell), rat IgG1 (clone HRPN, BioXcell), and mouse IgG2a (clone C1.18.4, BioXcell), were administered at the same dose and schedule. To deplete alveolar macrophages (AMs), mice received intranasal (*i.n.*) delivery of 50 µL clodronate liposomes or phosphate-buffered saline (PBS) liposome controls (Liposoma BV, The Netherlands) three times per week.

### Lung metastasis visualization and histology analysis

For metastasis visualization, mice were euthanized by carbon dioxide between days 25 and 34 after tumor inoculation or as otherwise indicated. Lungs were inflated with 15% Indian ink via intratracheal injection, then fixed overnight in Fekete’s solution (70% ethanol, 3.7% paraformaldehyde, 4% glacial acetic acid) to remove excess stain. After fixation, metastasis nodules appeared white and were counted under a Leica dissecting microscope.

For histology analysis, the lungs were perfused with PBS, fixed in 10% neutral buffered formalin overnight, processed for paraffin embedding, and sectioned at 5 µm for hematoxylin and eosin staining (H&E) or Immunohistochemistry (IHC). IHC was performed in-house using the Leica Bond RX Automated IHC Platform with anti-CD68 antibody (1:250, ab125212, Abcam), ER1 antigen retrieval solution (AR9961, Leica Biosystems), and BOND IHC Polymer Refine Detection System (DS9800, Leica Biosystems), followed by hematoxylin counterstaining. Slides were mounted and scanned using a Zeiss Axioscan 7 whole-slide scanner. Tumor area and macrophage infiltration were quantified using the HALO and HALO AI Image Analysis Platform (Indica Labs).

### Flow cytometry analysis of lung immune cells

To profile lung immune cells during metastasis progression, CMT167-bearing C57BL/6 mice were euthanized by CO₂ on days 7, 21, and 28 post tumor implantation, and 4T1-bearing BALB/c mice were euthanized on days 7, 14, and 28. For evaluation of macrophage depletion efficiency, mice were euthanized the next day following the third intranasal administration of clodronate or PBS liposomes, or following the seventh intraperitoneal dose of anti-CSF1R or isotype control antibody. For these depletion studies, intravascular leukocytes were labeled by retro-orbital injection of 2.5 µg of BB700 anti-CD45 antibody (BD Biosciences) in 100 µL of sterile PBS under isoflurane anesthesia. The antibody was allowed to circulate for 5-10 minutes before euthanasia (83).

Immediately after euthanasia, the lungs were perfused with PBS, harvested, and minced with scissors. The tissues were digested for 45 minutes at 37°C in complete RPMI-1640 supplemented with collagenase type IV (2 mg/mL; Gibco) and DNase I (20 µg/mL; Roche), as previously described (46). The digested tissues and cell suspension were passed through a 70 µm cell strainer and centrifuged at 400 g for 5 minutes to pellet the cells. Following red blood cell lysis with ACK buffer, the remaining cells were washed using PBS. For Fc receptor blocking, cells were incubated with 1 µg of anti-CD16/CD32 (BioXcell) in 100 µL of PBS per sample for 15 minutes at room temperature, in the presence of Zombie NIR viability dye (1:1000; BioLegend). Cells were then washed with FACS buffer (PBS without calcium and magnesium supplemented with 0.5% bovine serum albumin and 5 mM EDTA) and stained with the antibodies listed in Supplementary Table 2 for 45 minutes at 4°C. For intracellular staining, cells were fixed and permeabilized using the eBioscience™ Foxp3/Transcription Factor Staining Buffer Set (Invitrogen) according to the manufacturer’s instructions. Flow cytometry was performed on a Cytek Aurora (Cytek Biosciences), and data were analyzed using the FlowJo v10 software (Tree Star). For evaluation of macrophage depletion efficiency, the gating strategies for lung resident immune cell identification are provided in Supplementary Table 3.

### Western Blot

2.5 × 10⁵ cells per well were plated in 6-well plates, allowed to adhere, and treated as indicated overnight. Cells were harvested by scraping into 200 μL of RIPA lysis buffer (Thermo Fisher Scientific) supplemented with protease and phosphatase inhibitors (Thermo Fisher Scientific). Whole-cell lysates were prepared, and 20 μg of protein per sample was loaded on a 4-20% Criterion™ TGX™ Precast Midi Protein Gel (Bio-Rad), followed by transfer onto nitrocellulose membranes (Bio-Rad). Membranes were blocked with Intercept (TBS) Blocking Buffer (LI-COR) for 1 hour at room temperature, then cut by molecular weight range as appropriate. Immunoblotting was performed using primary antibodies against PTPN1 (Proteintech #11334-1-AP, RRID#AB_10642566), PTPN2 (Cell Signaling Technology #58935, RRID#AB_2799550), phospho-STAT1 (Cell Signaling Technology #9167, RRID#AB_561284), phospho-STAT3 (Cell Signaling Technology #9145, RRID#AB_2491009), and actin (Cell Signaling Technology #4970, RRID#AB_2223172) overnight at 4 °C. After washing in TBST (Tris-buffered saline with 0.1% Tween-20), membranes were incubated with fluorophore-conjugated secondary antibodies (LI-COR) and visualized using a LI-COR imaging system.

### Preparation of lung cell suspensions and library construction for scRNA-seq

BALB/c mice implanted with 4T1 tumors were treated daily with AC484 or vehicle control, starting on Day 7 after tumor inoculation. On Day 14, single-cell suspensions were prepared from PBS-perfused lungs as described above. The cells were stained with NucBlue Fixed Cell Stain (Invitrogen), and viable cells (NucBlue negative) were sorted using the FACSAria cell sorter (100 µm nozzle; BD Biosciences). Droplet-based single-cell capture and library preparation were conducted on a 10x Genomics Chromium Controller using the Chromium Next GEM Single Cell 5′ Reagent Kit v2.2, following the manufacturer’s protocol. Three lungs per treatment group were collected for this analysis.

### scRNA-seq pre-processing, quality control and cell type annotation

Raw FASTQ files were processed by Cell Ranger v9.0.0 (10x Genomics) using mm10-2020 as a reference. Cell Ranger aggregate was used to perform read-depth normalization. Cells expressing over 5% of mitochondrial transcripts or fewer than 100 total transcripts were excluded. Cells with transcript counts greater than the 95^th^ percentile within each sequencing experiment were considered doublets and therefore excluded. If not indicated otherwise, log-transformed normalized counts per ten-thousand (CP10K) were used throughout the study and the final data set was subset to highly variable genes based on a minimum mean of 0.0125, a maximum mean of 3, and a minimum dispersion of 0. These data were used for cell type annotation involving a combination of supervised and unsupervised methods. First, we applied SCANVI using a previously reported longitudinal scRNA-seq study in 4T1 as the reference data set (31). The assigned cell type labels by this classifier were then carefully assessed by computing differential expression analysis and manually benchmarking the results to established cell type markers from the literature and the CZ CELLxGENE Tabula Muris Senis atlas (41). All scRNA-seq analyses were performed with Scanpy unless otherwise specified. Rank GSEA was performed using the *prerank* function of gseapy version v1.1.10 with the gene set libraries GO, KEGG, and Oncogenic signature gene sets (C6, MSigDB). We performed differential expression analysis in Scanpy with the *sc.tl.rank_genes_groups* function, applying the Wilcoxon test.

### Dimensionality reduction, RNA velocity analysis, and clustering

In all embeddings, vehicle and AC484-treated samples were combined in the same object for comparison and subjected to dimensionality reduction using PCA (arpack svd solver) and UMAP using 30 principal components and 30 neighbors. For the analysis of neutrophils shown in Supplementary Figure 3C-E, Leiden clustering was performed using a resolution weight of 0.50. SCVELO was used with default parameters assuming the dynamical model of transcriptional dynamics to estimate RNA velocities (84,85). To orthogonally infer the differentiation trajectory of neutrophils, we also used an established signature of early neutrophil development (86). Neutrophil-specific transcriptional signatures associated with degranulation, cytotoxicity, ISG, tumor proliferation, and myeloid cell recruitment were obtained from Gungabeesoon et al (39), whereas the Ly6E^hi^ signature was retrieved from Benguigui et al (40). To identify differentially-abundant cell clusters specific to vehicle and AC484-treated animals, we selected either neutrophils only or the rest of the cells and used them as inputs for DAseq in R (38). For DAseq clustering of neutrophils, we used a vector of k weights spanning 50 to 1000 in steps of 50 to calculate the DA scores for each cell. We set the top and bottom DA score threshold in -0.85 and 0.85, respectively, to exclusively focus on cells that were most specific to each treatment group and filter out the rest. We used a resolution of 0.05 for the identification of DA regions in this model. To run DAseq on the rest of the cell types, excluding neutrophils, we used the same parameter but relaxed the DA score thresholds to -0.52 and 0.52. The DAseq STG_markers function was used to identify markers for the generated clusters. To characterize the phenotype of alveolar macrophages, we used functional signatures reported in a previously published scRNA-seq study analyzing 4T1 lung metastasis (41).

### Spatial transcriptomics sample preparation and data acquisition

BALB/c mice bearing 4T1 tumors were treated daily with AC484 or a vehicle control starting Day 7 post-tumor inoculation. Mice were humanely euthanized by CO_2_ on Day 22. Immediately following euthanasia, the lungs were perfused with PBS, harvested, and fixed in 10% neutral buffered formalin overnight. Tissues were then processed for paraffin embedding and sectioned at 5 µm and placed on Superfrost Plus slides (Fisherbrand, 22-037-246). All procedures were conducted in an RNase-free environment.

Spatial transcriptomics was performed on the slide using the CosMx Mouse Universal Cell Characterization RNA Panel according to the manufacturer’s protocol: CosMx SMI Manual Slide Preparation for RNA Assays (FFPE; MAN-10184-04). Briefly, the slide was placed in a 60°C incubator for a 2-hour bake. The tissue sections were then deparaffinized with xylene and washed with ethanol. The slide was rehydrated and underwent target retrieval in a pressure cooker. The slides were allowed to dry for at least 30 minutes but no longer than 1 hour. The tissue sections were permeabilized with a combination of proteinase K treatment following the dilution ratio listed in the manual. The fiducials were then prepared and applied to the slides. After application of the fiducials, the tissue sections were fixed with 10% NBF, blocked, and incubated overnight at 37°C in an ACD hybridization oven (PN 321710/321720) with denatured CosMx Mouse Universal RNA panel, 1000-plex, without add-on probes. The following day, within 18 hours, the slides were washed, blocked, and stained with CosMx Mouse Universal Cell Segmentation Kit (RNA), including nuclear (DAPI) and cell segmentation markers (CD298/B2M and PanCK/CD45). Prior to loading the CosMx instrument, the CosMx RNA imaging tray 1000-plex was allowed to come to room temperature for at least 1 hour, and RNase Inhibitor was added immediately before loading of the imaging tray. The slides were loaded into the CosMx instrument, inverted with a pre-bleaching profile of Configuration C and a cell segmentation profile of Configuration A. With all reagents loaded following the loading manual (MAN-10161-10), the slides were scanned, and the fields of view were selected to cover the entire section or regions of interest for RNA profiling.

### Spatial transcriptomics data analysis

Assembly and segmentation: The CosMx section was segmented in AtoMx using Config A - Non-Neuro Human Tissue and systematically evaluated for select FOVs. Raw files were then exported from the AtoMx platform and analysis was applied in R 4.5.1 using R scripts from Nanostring (https://github.com/Nanostring-Biostats) and custom scripts as described below.

Preprocessing and quality control: Raw CosMx data, including cell segmentation masks, transcript locations, and cell-by-gene expression matrices, were loaded for analysis. The initial dataset was organized by tissue type and field of view (FOV). Quality control was performed to remove low-quality cells and FOVs according to Nanostring’s recommendations. Cells with fewer than 100 counts were also excluded from further analysis. Gene expression data was normalized to probe detection depth for each cell.

Unsupervised clustering: Clustering was performed using Nanostring’s *InSituType* approach. Briefly, cohorts were constructed using mean fluorescent signal from each marker in the CosMx panel, and raw probe counts, per-cell background signal, and mean negative probe signal were supplied. Clusters were evaluated according to separation in higher dimensional space and distinction in marker gene expression to iteratively merge and subcluster using the refineClusters function.

Cell type annotation: Initial cell type annotation was performed on the unsupervised clusters by examining the expression of canonical marker genes. Differential expression analysis between clusters using FindAllMarkers in Seurat (Wilcoxon rank-sum test) was employed to identify cluster-specific markers. These markers, in conjunction with known lineage-specific genes, were used to assign biological labels to each cluster. Cell type labels were further refined and finalized through iterative analysis and comparison with reference atlases.

Spatial analysis: The spatial coordinates of each cell were used to analyze the spatial organization of cell types. To investigate the tumor microenvironment, the Euclidean distance from each cell to the nearest tumor cell was calculated. Based on this distance (∼300 μm), cells were categorized into discrete proximity groups (“proximal”, “distal”).

Differential expression and gene set enrichment analysis: Differential gene expression (DGE) analysis was conducted to compare cell states between experimental conditions (AC484 vs. Vehicle) within specific cell types and spatial contexts (e.g., tumor-proximal versus -distal alveolar macrophages). The FindMarkers function in Seurat was used for this purpose. Genes were considered significantly differentially expressed with an adjusted p-value < 0.05. Gene Set Enrichment Analysis (GSEA) and Over-Representation Analysis (ORA) were performed on the lists of differentially expressed genes to identify enriched biological pathways and functions. The clusterProfiler and enrichR packages were used, with gene sets from the MSigDB Hallmark (H) and Gene Ontology (C5) collections.

Cell-cell communication analysis: To infer intercellular communication networks, CellChat was employed. The analysis was run separately on data from each experimental condition. Communication probabilities between all pairs of cell types were calculated based on the expression of known ligand-receptor pairs. The strength of communication pathways was compared between conditions to identify treatment-induced changes in cellular crosstalk.

### Multiplex immunofluorescence (IF) to quantify AMs in the lung

Mice were humanely euthanized using CO_2_ at Day 16 after tumor inoculation as indicated. Immediately following euthanasia, the lungs were perfused with PBS, harvested, and fixed in 10% neutral buffered formalin overnight. Tissues were then processed for paraffin embedding.

Formalin-fixed and paraffin-embedded lung tissues were sectioned to a thickness of 5 µm and mounted on glass slides. Sections were de-paraffinized, rehydrated, and antigen retrieval was performed using a steamer, by heating the antigen retrieval buffer (Biocare Medical, DV2004MM) to 95°C and steaming the slides for 20 minutes. After cooling, samples were placed in a Sequenza rack (Fisher Scientific, 73310017) and permeabilized in 0.05% Triton™ X-100 (Sigma, T8787) in PBS for 10 minutes at room temperature. Samples were then placed in blocking buffer, 5% BSA (Sigma, A2153) in PBS-T (PBS with 0.01% Tween 20, Sigma, P9416), to reduce non-specific binding for 1 hour at room temperature. Primary antibodies: rabbit anti-CD68 (1:100, Abcam, ab125212) and rabbit anti-CD11b (1:100, Abcam, ab133357) were conjugated to CoraLite® 647 and 750 dyes, respectively, using FlexAble 2.0 Antibody Labeling Kits (Proteintech Group, Inc.) according to the manufacturer’s protocols. Additionally, Hydrazide-AF488 (1:500 dilution, AAT bioquest, 1364) was added to the primary cocktail. The resulting fluorescently labeled primary antibody cocktail was diluted in blocking buffer, and incubated with the samples overnight at 4°C. Following primary antibody incubation, samples were washed three times for 5 minutes each in PBS-T at room temperature. Samples were then counterstained with Hoechst 33342 (1:1000, Invitrogen, H3570) in PBS for 10 minutes at room temperature. Slides were finally washed three times in PBS-T for 5 minutes each, gently dried, and mounted using ProLong Glass Antifade Mountant (ThermoFisher, P36980) with #1.5 coverslips (Marienfeld, Germany).

Fluorescence images were captured with an Axioscan 7 Slide Scanner (Zeiss, Germany). Images were acquired using a Plan-Apochromat 20x/.8 M27 objective, and the following acquisition channels: 430 nm, 475 nm, 630 nm and 750 nm, for Hoechst, Hydrazide, CoraLite®-647 and 750nm dyes, respectively. Data acquisition was managed through the Zeiss ZEN software user interface (Zeiss, Germany).

### Whole-slide multiple IF image analysis

Whole-slide multiplex IF of lung sections (DAPI-AF405, Hydrazide AF488, CD68-AF647, CD11b-AF750) were run in arivis Vision4D (ZEISS arivis Pro 4.3.1) using custom pipelines and models. Briefly, to generate the tissue mask, a 2-channel pixel classifier (inputs: CD68 and DAPI) separated tissue from background/blood. Objects <50 µm^2^ were removed. This mask constrained all downstream steps. To generate tumor segmentations, 2-channel DL model (inputs: DAPI and CD11b) segmented tumor vs background. Masks were regularized by filling inclusions, 2D opening (sphere, 15 px), and merging. Tissue gaps <1,000 µm^2^ were closed, then the tumor was intersected with the gap-filled tissue to enforce boundaries. Conducting airways were segmented with a separate DL model (inputs: Hydrazide and DAPI) and subtracted from tumor. Final tumors required area >=1,000 µm^2^. Tumors were binned as micro (<80,000 µm^2^) or macro (>=80,000 µm^2^).

For cell segmentation, nuclei were segmented with Cellpose (model “nuclei”, diameter 8.25 µm, 1-99% normalization, min area 5 µm^2^) on DAPI. Nuclei were expanded to whole-cell objects by watershed region growing on CD11b (AF750; threshold 400; max distance 87.6 µm). Cells intersecting airway ROIs were removed. Size gating kept 25-600 µm^2^ projected area. For signal correction, CD68 and CD11b channels were denoised (discrete Gaussian, 1.03 µm) and background-corrected (morphological top-hat, radius 2.75 µm, sphere). To determine macrophage subtype, per-cell mean intensities were measured on background-corrected CD68 and CD11b. Positivity thresholds were set per slide from intensity histograms (global thresholding). Macrophages were defined as AM = CD68^+^/CD11b^-^ and IM = CD68^+^/CD11b^+^. To generate compartment-specific metrics, the tissue was partitioned into Tumor (union of all tumors) and Tumor-free (tissue minus tumor). For each compartment, we reported densities for total CD68^+^ macrophages, AMs, and IM. Three independent lungs per treatment group were analyzed.

### Cytokine analysis of serum and bronchoalveolar lavage fluid (BALF)

For experiments where only serum was analyzed, mice were humanely euthanized by CO₂. Blood was collected via cardiac puncture into serum separator tubes, allowed to clot for around 45 min, and then centrifuged at 2,000 g for 15 min. The resulting supernatant as serum was analyzed using the Olink Target 48 Mouse panel.

For experiments requiring both serum and BALF, mice were euthanized via an intraperitoneal (i.p.) injection of 200 mg/kg Euthasol. Blood was first collected as described above. Two sequential bronchoalveolar lavage (BAL) samples were then collected from each mouse via the trachea using 0.75 mL of 2 mM EDTA-HBSS. The pooled BAL samples were centrifuged at 400g for 5 min at 4°C, and the supernatant was collected as BALF. Prior to analysis, BALF was concentrated five-fold using a 3 kDa MWCO Amicon Ultra Centrifugal Filter (Millipore). Cytokine analysis of both serum and BALF was performed with the LEGENDplex Mouse Cytokine Release Syndrome Panel (13-plex) (BioLegend) according to the manufacturer’s instructions. The panel measured: IFNγ, IL-10, CCL4, IFN-α, CXCL9, CXCL10, TNF-α, IL-6, VEGF, IL-4, CCL3, CCL2, and GM-CSF. LEGENDplex Data Analysis software (BioLegend) was used to determine cytokine concentrations based on standard curves.

### *In vitro* characterization and stimulation assays on BAL AMs

Mice were euthanized by i.p. injection of Euthasol (200 mg/kg) about 2 hours after the final dose of AC484 or vehicle control, between days 14-27 post-tumor implantation, with treatment initiated on day 7. Two bronchoalveolar lavages were performed on each mouse via the trachea using 0.75 mL of 2 mM EDTA-HBSS per lavage. The samples were then pooled to generate a single-cell suspension and centrifuged at 400 g for 5 min at 4 °C to pellet the cells.

For intracellular staining of IFNγ and CXCL9 in AMs, BAL cells from three mice were pooled, washed in RPMI, resuspended in complete RPMI medium, and seeded into ultra-low attachment 96-well plates (Corning, CLS4520). After overnight incubation at 37°C, cells were treated with Brefeldin A (5 μg/mL; BioLegend) and Monensin (2 μM; BioLegend) for 4 h, then harvested, fixed, and permeabilized using the Cyto-Fast Fix/Perm Buffer Set (BioLegend). Staining was performed with BV750 anti-CD11c (Clone N418, BioLegend), Kiravia Blue 520 anti-CD170 (Clone S17007L, BioLegend), APC anti-IFNγ (Clone XMG1.2, BioLegend), and PE anti-CXCL9 (Clone MIG-2F5.5, BioLegend). AMs were identified as CD11c⁺/CD170⁺.

For interferon stimulation, BAL cells from individual mice were divided into three equal fractions, resuspended in Hanks’ Balanced Salt Solution (with calcium/magnesium, Gibco), and incubated at 37°C for 15 min with recombinant mouse IFNγ (150 ng/mL; PeproTech, 315-05), IFNα (150 ng/mL; BioLegend), or an equivalent volume of vehicle control (0.5% BSA). Cells were then immediately fixed in 4% paraformaldehyde for 10 min at 37°C, permeabilized with cold Perm III buffer (BD Biosciences) on ice, and stained with BUV496 anti-CD45 (Clone 30-F11, BD Biosciences), Kiravia Blue 520 anti-CD170 (Clone S17007L, BioLegend), and Alexa Fluor 647 anti-STAT1 pY701 (Clone 4a, BD Biosciences). AMs were identified as CD45⁺/CD170⁺. The level of p-STAT1 was calculated as the fold change in geometric mean fluorescence intensity (gMFI) relative to the vehicle control (unstimulated AMs).

To examine p-STAT1 after LPS stimulation, BAL cells from individual mice were resuspended in complete RPMI, split into two groups, and seeded in ultra-low attachment 96-well plates. One group was stimulated with 2.5 µg/mL LPS (eBioscience) for 2.5 h, while the other group was left untreated, followed by immediate fixation and staining for p-STAT1. The gMFI of p-STAT1 was normalized to the untreated controls to determine fold change. For chemokine analysis, BAL cells pooled from three mice per group were stimulated with LPS for 2.5-3 h in complete RPMI, then treated with Brefeldin A (5 μg/mL; BioLegend) and Monensin (2 μM; BioLegend) overnight. Cells were then fixed and permeabilized with the Cyto-Fast Fix/Perm Buffer Set (BioLegend) and stained with BV750 anti-CD11c, Kiravia Blue 520 anti-CD170, and PE anti-CXCL9. AMs were identified as CD11c⁺/CD170⁺.

All representative data were obtained from at least two independent experiments or more than three biological replicates, with each mouse considered a single replicate.

### *In vitro* evaluation of p-STAT1 in AMs cocultured with tumor cells

Mice bearing 4T1 or CMT167 tumors were used to generate AMs for matched tumor cell types for *in vitro* assays. Animals were euthanized with Euthasol (200 mg/kg) between days 14-30 post-tumor implantation, and bronchoalveolar lavage (BAL) cells were harvested as previously described.

To assess p-STAT1 activation in AMs, BAL cells from individual mice were resuspended in complete RPMI medium, divided into three groups, and seeded in ultra-low attachment 96-well plates. The BAL cells were then co-cultured in three groups: 1) 4T1 or CMT167 tumor cells and AMs treated with AC484 (100 nM); 2) tumor cells and AMs treated with DMSO vehicle; 3) a control group consisting of AMs cultured alone with DMSO. AMs cocultured with 4T1 were collected after overnight incubation, whereas coculture with CMT167 was maintained for 48 hours. Following coculture, cells were fixed with 4% paraformaldehyde, permeabilized with cold Perm III buffer, and stained with BV750 anti-CD11c, Kiravia Blue 520 anti-CD170, and Alexa Fluor 647 anti-STAT1 pY701 as previously described. AMs were identified as CD11c⁺/CD170⁺. The gMFI of p-STAT1 was normalized to the unstimulated control group (AMs cultured alone with DMSO vehicle) to determine fold change.

### AM-mediated tumor killing assay

As previously described, BAL cells were harvested from tumor-bearing mice. AMs were selectively counted based on their larger size using a hemocytometer. 5 x 10^4^ AMs per well were seeded into black-wall 96-well plates containing complete RPMI. After a 2 h incubation at 37 °C, non-adherent cells were removed by two gentle PBS washes, leaving adherent AMs. Tumor cells (1.25 × 10³ 4T1-Luc or CMT167-Luc) were then added, followed by recombinant mouse IFNγ (PeproTech) at the indicated concentrations or vehicle control (0.5% BSA), along with AC484 (300, 100 or 50 nM, as indicated) or DMSO. The final AM-to-tumor cell ratio was 40:1, with a total culture volume of 150 μL per well.

In experiments involving the JAK inhibitor ruxolitinib, BAL cells were pre-incubated with ruxolitinib (3 μM, MedChemExpress) in complete RPMI for 2 h, washed, and then cocultured as described, with ruxolitinib maintained throughout the assay at 3 μM. Equivalent DMSO volumes were used as vehicle controls.

For transwell assays, 1.0 μm pore inserts (Corning, CLS3380) were pre-equilibrated overnight in complete RPMI. BAL AMs (5 × 10⁴ per well) were seeded onto the inserts, incubated for 2 h, and gently washed twice using PBS. Tumor cells were seeded into black-wall 96-well plates, and inserts containing AMs were placed above tumor cells. Cultures were supplemented with IFNγ (1 ng/mL) and AC484 (150 nM) or vehicle controls from the top of the insert. AMs seeded in contact with tumor wells served as controls.

Tumor cell viability was assessed 3 days after coculture using the ONE-Glo Luciferase Assay System (Promega) according to the manufacturer’s instructions. Briefly, the reagent was added to each well at a 1:1 ratio with the culture medium, plates were shaken at 600 rpm for 3 min, and luminescence was measured using a Victor X2 Microplate Reader (PerkinElmer). The luminescence readout from 4T1-Luc or CMT167-Luc cells cultured under unmanipulated control conditions (no AMs or AC484 added) was defined as 100% viability and used to normalize the data.

For cytokine and chemokine measurements, 100 μL of coculture supernatant was collected, centrifuged at 400 × g for 5 min, and supernatants were analyzed using the LEGENDplex Mouse Cytokine Release Syndrome Panel (13-plex; BioLegend) following the manufacturer’s instructions as described before.

### Statistical Analysis

All statistical analyses were conducted in GraphPad Prism. We used an ANOVA test for comparing more than two groups and an unpaired t-test for two-group comparisons. Statistical significance was defined as a p-value ≤ 0.05. Resulting p-values are denoted as p > 0.05 (ns), p ≤ 0.05 (*), p ≤ 0.01 (**), and p ≤ 0.001 (***).

### Data Availability

All data generated in this study are included in the article and its supplementary files. The single-cell RNA sequencing, spatial transcriptomics, and bulk RNA sequencing data will be deposited in the Gene Expression Omnibus under accession number upon publication. Additional information is available from the corresponding author upon reasonable request.

## Author contributions

Conceptualization: Y. Liu, J. D. Powell

Methodology: Y. Liu, I.-M. Sun, M. Creixell, J. Brown, S. Kharbanda, J. J. Lee, V. Shahryari, K. Hake, J. O’Hara, N. Yang, L. Penland, J. Wang, K. M. Li, J. Balibalos, A. W. Stebbins, P. M. Godfrey, P.-H. Tai, E. Malahias, W. Kong, N. Fong, L. Chan, C. H. Patel, T. A. Nguyen

Formal analysis: Y. Liu, I.-M. Sun, M. Creixell, J. Brown, S. Kharbanda, J. J. Lee, V. Shahryari, K. Hake, S. Gupta, T. A. Nguyen

Data curation: Y. Liu, I.-M. Sun, M. Creixell

Supervision: J. J. Lee, K. J. Finn, D. Hendrickson, F. McAllister, M. N. Paddock, T. A. Nguyen, F. A. Harding, J. D. Powell

Writing - original draft: Y. Liu, I.-M. Sun, M. Creixell, T. A. Nguyen, F. A. Harding, J. D. Powell

Writing - reviewing & editing: Y. Liu, I.-M. Sun, M. Creixell, J. Brown, S. Kharbanda, J. J. Lee, V. Shahryari, K. Hake, J. O’Hara, K. J. Finn, N. Yang, L. Penland, J. Wang, K. M. Li, J. Balibalos, A. W. Stebbins, P. M. Godfrey, P.-H. Tai, E. Malahias, W. Kong, N. Fong, D. Hendrickson, S. Gupta, L. Chan, F. McAllister, C. H. Patel, M. N. Paddock, T. A. Nguyen, F. A. Harding, J. D. Powell

## Acknowledgments

We thank the Laboratory Animal Resources department (Ellie Karlsson, Karla Barron, Marcelo Cosino, Raul Garcia-Gonzalez, Ronny Luna, Samira Nazertehrani, Paulyn Cha, and Natalie Smolnikov) for vivarium support, and Amy Jo Johnson for coordinating genomic sequencing services. We are grateful to our colleagues at AbbVie for their valuable discussions and for generously providing AC484 and vehicle. We thank Magdalena Preciado López, Austin Lefebvre, and Anatalia Robles for reviewing the manuscript. We also appreciate the support of our Calico colleagues, particularly Wen Lu, Agnieszka Anya Wendorff, Alexander Li, and Astrid Gillich for insightful discussions, and David Stokoe and Cheng-yen Chang for their critical review and expert advice.

We acknowledge the use of Gemini (Google) to assist with editing and improving the readability of the manuscript text. We reviewed all AI-suggested changes and retain full responsibility for the content. Figures were created with BioRender.com as indicated.

