## Supplementary Figure 1-11, Supplementary Table 1-3 for "PTPN1/2 inhibits alveolar macrophage-mediated control of lung metastasis"

### Figure S1, related to Figure 1

A

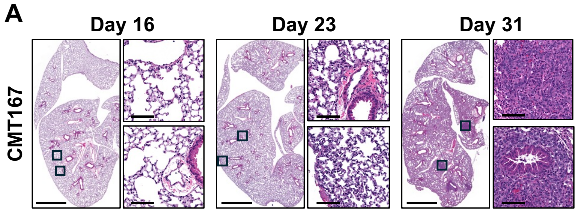


B

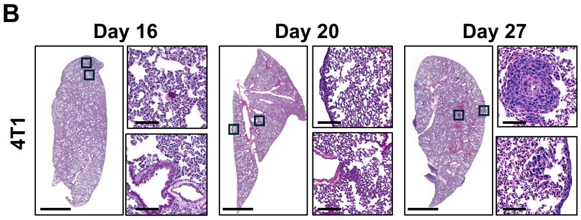


C Lung metastatic tumor area over time (CMT167 model)

| | Day 16 | Day 23 | Day 31 |
|---|---|---|---|
| Mean | 0.010 | 0.015 | 2.060 |
| SEM | 0.000 | 0.015 | 0.771 |

D Lung metastatic tumor area over time (4T1 model)

| | Day 16 | Day 20 | Day 27 |
|---|---|---|---|
| Mean | 0.064 | 0.645 | 7.403 |
| SEM | 0.000 | 0.000 | 4.741 |

E

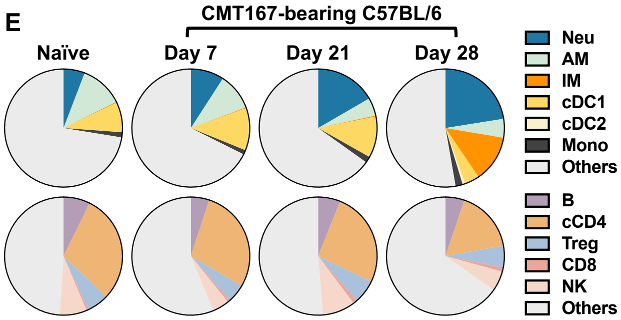


G

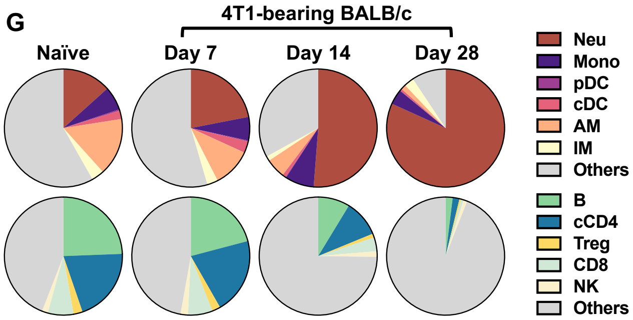


F

Immune cell % over time in the lung ( myeloid panel in CMT167 model)

| | Naïve | Day 7 | Day 21 | Day 28 |
|---|---|---|---|---|
| Neu | 5.836 | 9.156 | 16.65 | 21.12 |
| AM | 11.81 | 10.138 | 4.9375 | 4.846 |
| IM | 0.25 | 0.202 | 0.28 | 12.09 |
| Mono | 8.158 | 11.58 | 11.26 | 3.814 |
| cDC1 | 0.0666 | 0.0496 | 0.03225 | 0.654 |
| cDC2 | 1.408 | 1.21 | 1.685 | 1.84 |
| Others | 72.4066 | 67.6122 | 65.136 | 49.552 |

Immune cell % over time in the lung ( lymphocyte panel in CMT167 model)

| | Naïve | Day 7 | Day 21 | Day 28 |
|---|---|---|---|---|
| NK | 7.282 | 5.046 | 6.0625 | 5.214 |
| B | 29.82 | 28.38 | 26.05 | 16.96 |
| cCD4 | 5.988 | 5.042 | 6.565 | 5.958 |
| Treg | 0.532 | 0.68 | 1.035 | 1.134 |
| CD8 | 7.432 | 4.67 | 8.9825 | 5.496 |
| Others | 48.946 | 56.182 | 51.305 | 65.238 |

H

Immune cell % over time in the lung ( myeloid panel in 4T1 model)

| | Naïve | Day 7 | Day 14 | Day 28 |
|---|---|---|---|---|
| Neu | 13.35 | 21.95 | 51.23 | 81.78 |
| Mono | 6.53 | 6.48 | 7.95 | 4.15 |
| pDC | 0.35 | 0.25 | 0.19 | 0.14 |
| cDC | 2.3 | 3.15 | 0.96 | 0.49 |
| AM | 15.8 | 10.65 | 5.37 | 1.39 |
| IM | 3.47 | 2.98 | 1.29 | 2.72 |
| Others | 58.22 | 54.56 | 33.01 | 9.35 |

Immune cell % over time in the lung (lymphocyte panel in 4T1 model)

| | Naïve | Day 7 | Day 14 | Day 28 |
|---|---|---|---|---|
| B | 24.4 | 20.9 | 8.71 | 1.94 |
| cCD4 | 20.33 | 20.85 | 10.1 | 1.86 |
| Treg | 2.51 | 2.28 | 1.07 | 0.45 |
| CD8 | 7.05 | 6.74 | 3.78 | 0.72 |
| NK | 1.61 | 2.05 | 1.57 | 0.99 |
| Others | 44.09 | 47.19 | 74.77 | 94.05 |

**Figure S1 (related to Figure 1). Characterization of CMT167 and 4T1 spontaneous metastasis model**

**A, B.** Representative H&E staining of lung sections showing metastasis progression in CMT167 (**A**) and 4T1 (**B**) models at the indicated time points. Scanned half-lung regions and magnified views are shown. Scale bars: 2 mm (overview) and 100 µm (magnification). **C-D**. Quantification of parenchymal metastases based on H&E-stained lung sections collected at the indicated time points after implantation of CMT167 lung cancer cells (**C**) and 4T1 breast cancer cells (**D**). Data represent the percentage of tumor regions relative to total scanned lung area. Three independent animals were analyzed per time point; data are shown as mean and SEM. **E-H.** Immune profiling of lung immune cell populations during metastasis in the CMT167 (**E and F**) and 4T1 (**G and H**) models. Lungs were perfused with PBS, harvested, dissociated, and stained for flow cytometry analysis. Data represent averages from ≥ 3 mice.

#### Figure S2, related to Figure 1

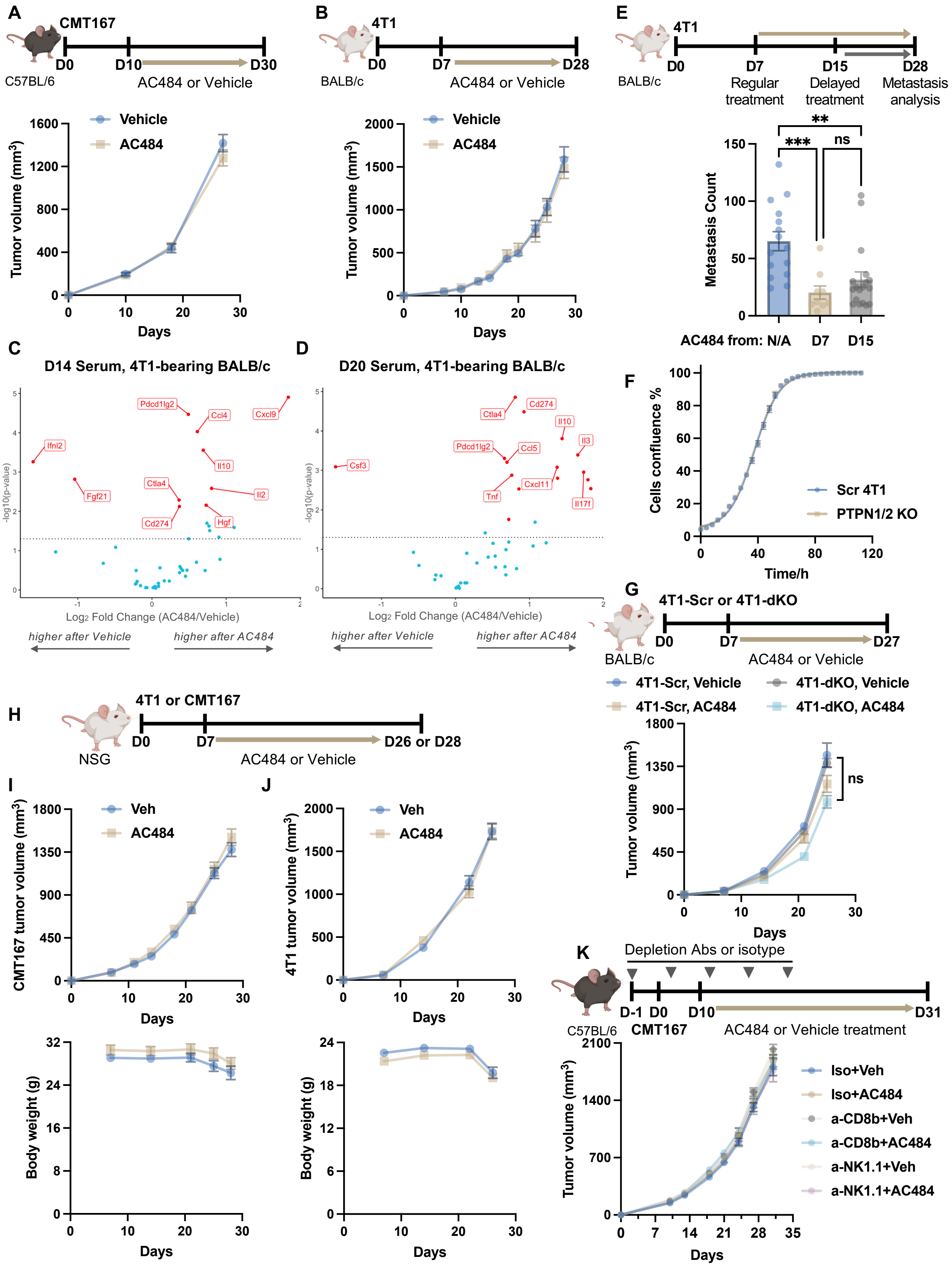

**Figure S2 (related to Figure 1). The anti-metastatic effects of AC484 are independent of its impact on primary tumor growth in CMT167 and 4T1 tumor models**

**A, B.** Experimental schedules as in Figure 1A-H and primary tumor growth kinetics of CMT167 (**A**) and 4T1 (**B**) tumors. **C, D.** comparison of serum cytokine levels in 4T1-bearing BALB/c mice treated with AC484 or vehicle control, collected on Day 14 (**C**) or Day 20 (**D**) post-implantation, with treatment starting on Day 7. Data represent the mean of N = 7 mice per group. **E.** BALB/c mice implanted with 4T1 cells received AC484 or vehicle control treatment starting on Day 7 or Day 15. Lung surface metastasis nodules were counted and shown as individual animals (dots) with mean ± SEM (bars). **F.** Growth of PTPN1/2 double knockout (dKO) and scrambled (Scr) control 4T1 cells *in vitro*. Cell confluence was quantified from ≥ 3 wells, shown as mean ± SEM (bars). **G.** Tumor growth kinetics of PTPN1/2 dKO and Scr 4T1 cells in BALB/c mice with the indicated treatment. **H-J.** Tumor growth kinetics of 4T1 (I) and CMT167 (J) tumors and corresponding body weight in NSG mice. **K.** Tumor growth kinetics of CMT167 tumors in C57BL/6 mice treated with the indicated depletion antibodies and compounds. For all tumor growth kinetics, shown are mean ± SEM (bars). N ≥ 7 mice per group. One-way ANOVA test: ns, $p > 0.05$; * $p \leq 0.05$; ** $p \leq 0.01$; *** $p \leq 0.001$.

#### Figure S3, related to Figure 2

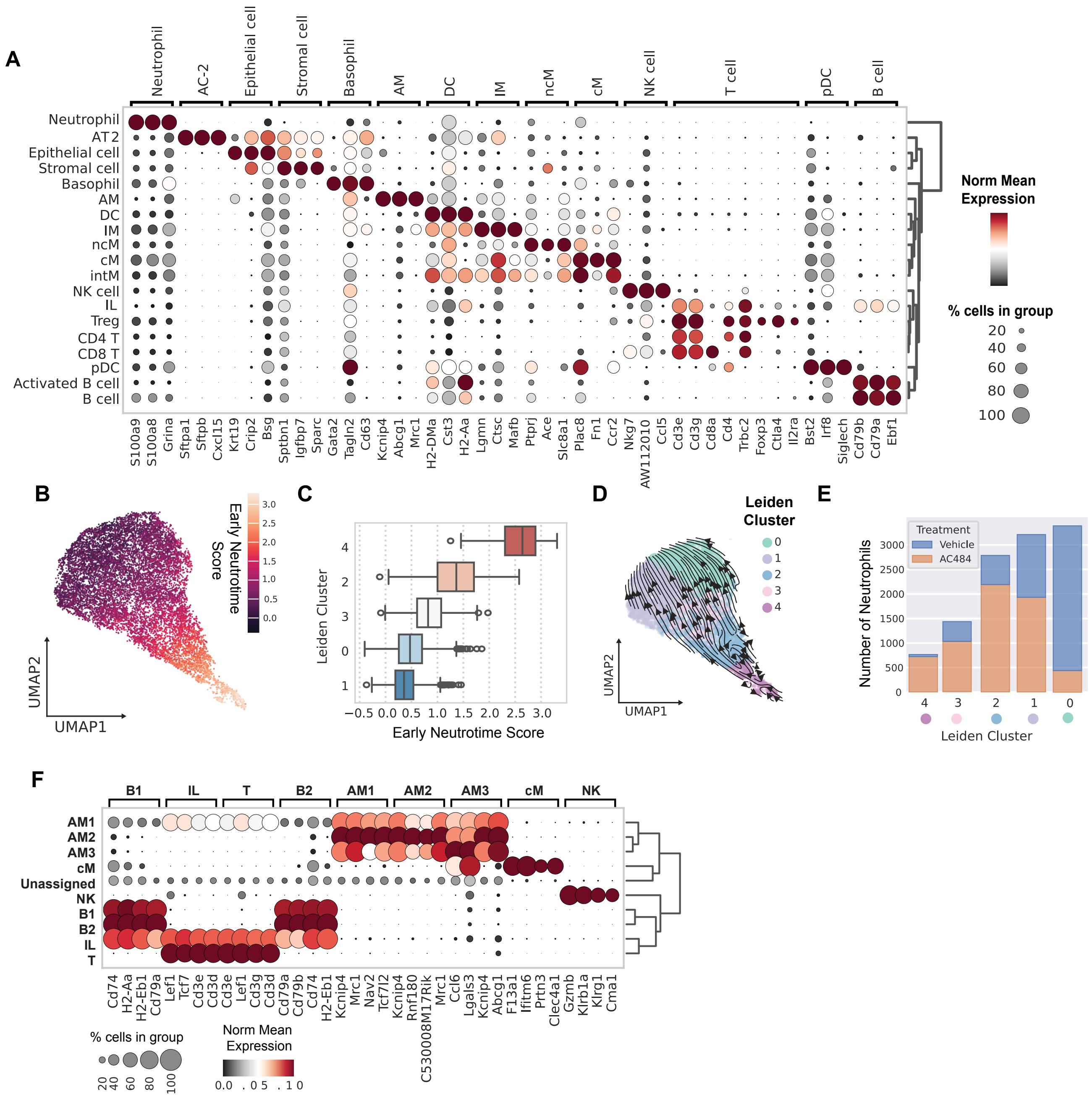


**Figure S3. (related to Figure 2). scRNA-seq markers and other supplementary plots**

**A**. Dot plot of markers for each cell type identified. **B-C**. Early neutrotime score of neutrophils. A higher value indicates a higher expression of genes involved in early neutrophil development. **D**. Trajectory analysis of neutrophils grouped by Leiden cluster. **E.** Bar plot showing the fraction of neutrophils grouped by treatment across Leiden clusters. **F.** DA subpopulation markers for non-neutrophil immune cell types.

**Figure S4, related to Figure 2**

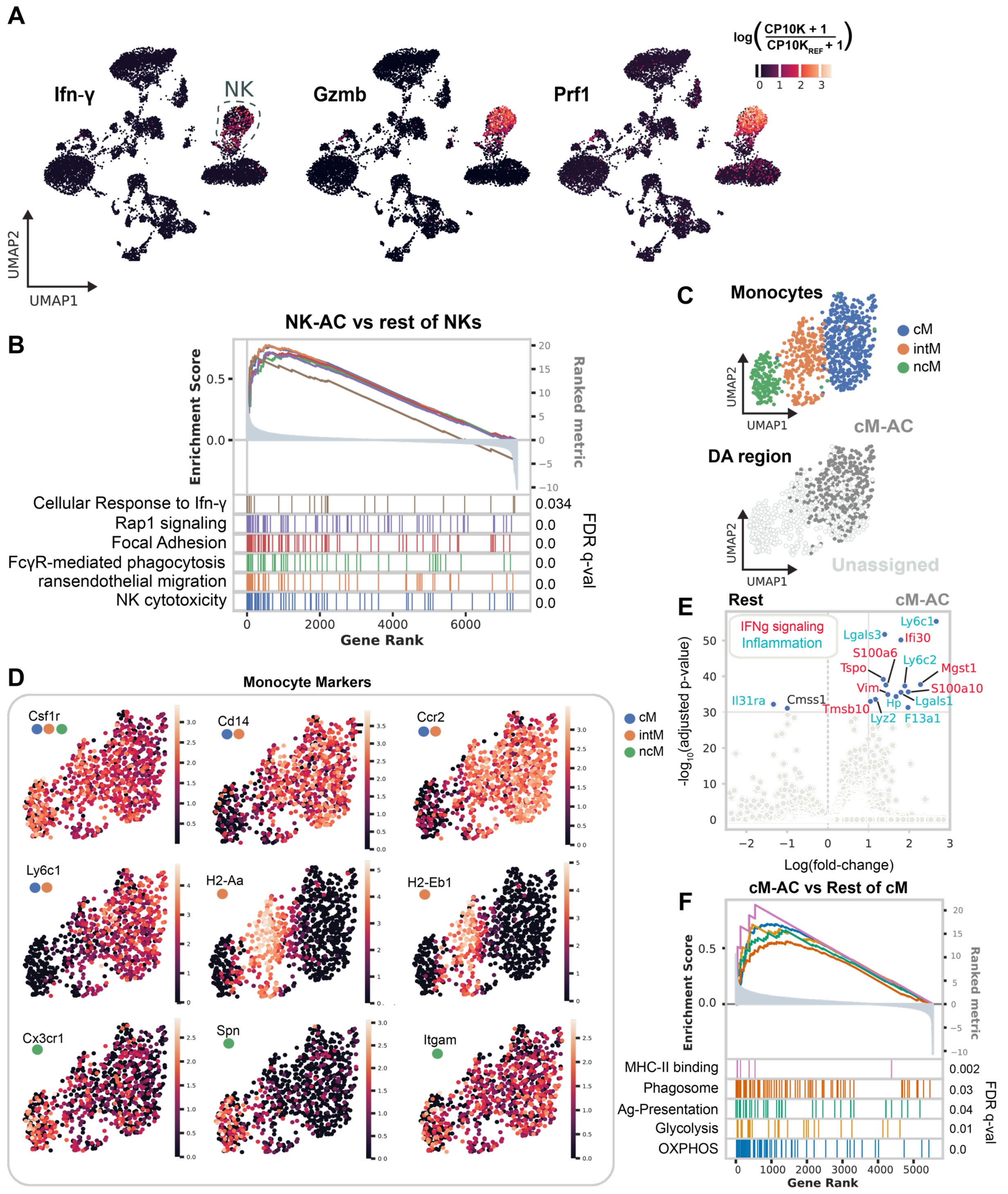


**Figure S4. (related to Figure 2). Analysis of additional DA subpopulations enriched in AC484-treated mice**

**A.** UMAP of all cells detected, excluding neutrophils and highlighting the expression of Ifng, Gzmb, and Prf1. **B.** GSEA analysis of NK-AC484 compared with the rest of NK in the dataset. **C**. UMAP of monocytes colored by classical (cM), intermediate (intM), or nonconventional (ncM) monocytes (top) and the same UMAP highlighting the DA subpopulation of cM enriched in AC484-treated mice (bottom). **D**. Markers defining cM, intM, and ncM. **E**. Differential expression analysis of cM-AC484 versus the rest of cM in the dataset, highlighting genes involved in IFNγ signaling and inflammation. **F.** GSEA of cM-AC484 compared with the rest of cM.

#### Figure S5, related to Figure 3

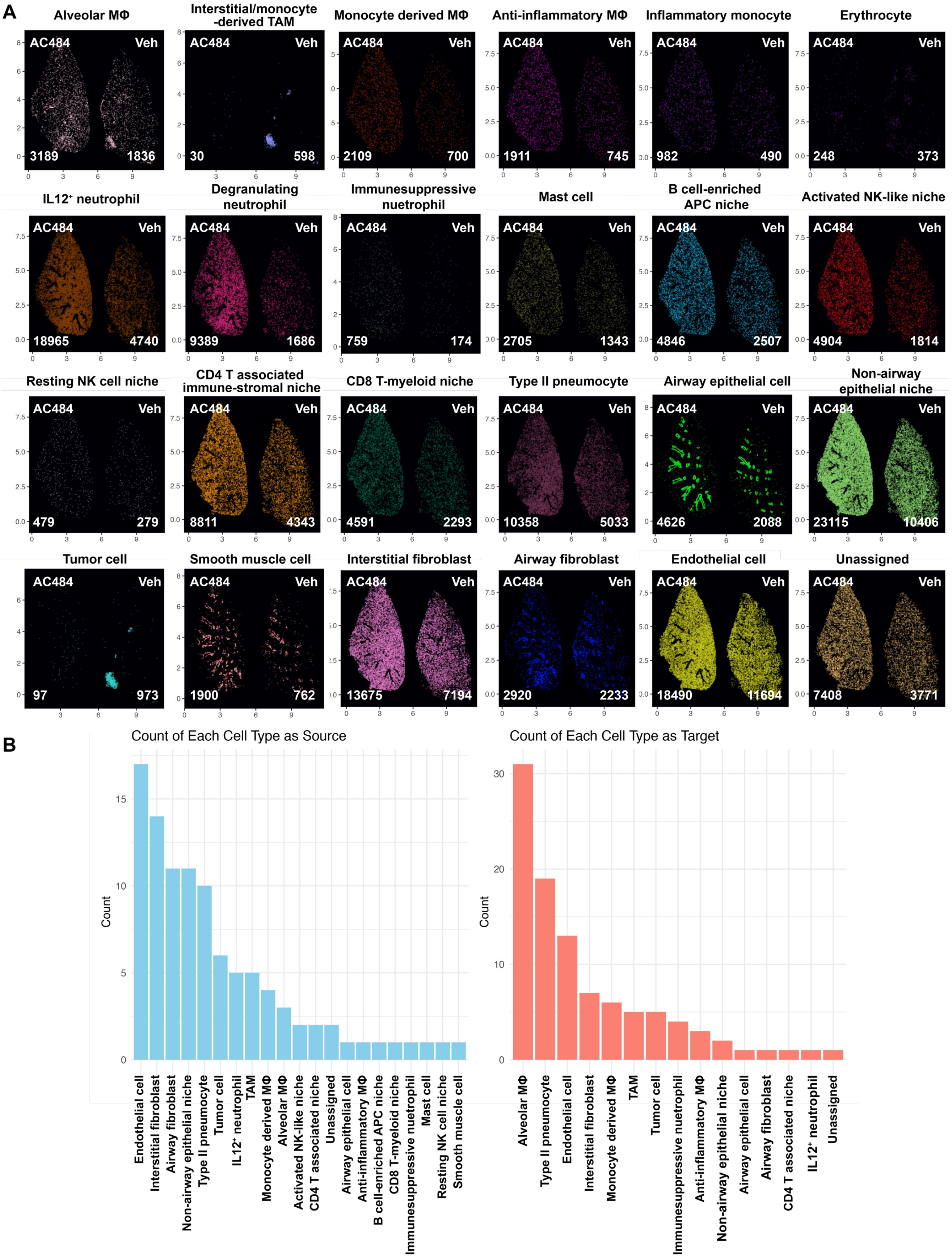

**Figure S5 (related to Figure 3). Spatial transcriptomics analysis identifies AMs as the predominant signal-receiving population**

**A**. Spatial maps and cell type annotations of the lung section in Figure 3C. Numbers indicate interaction counts per cell type. **B**. CellChat summary showing overall outgoing and incoming signal strength for each cell type, integrated across all pathways.

### Figure S6, related to Figure 3

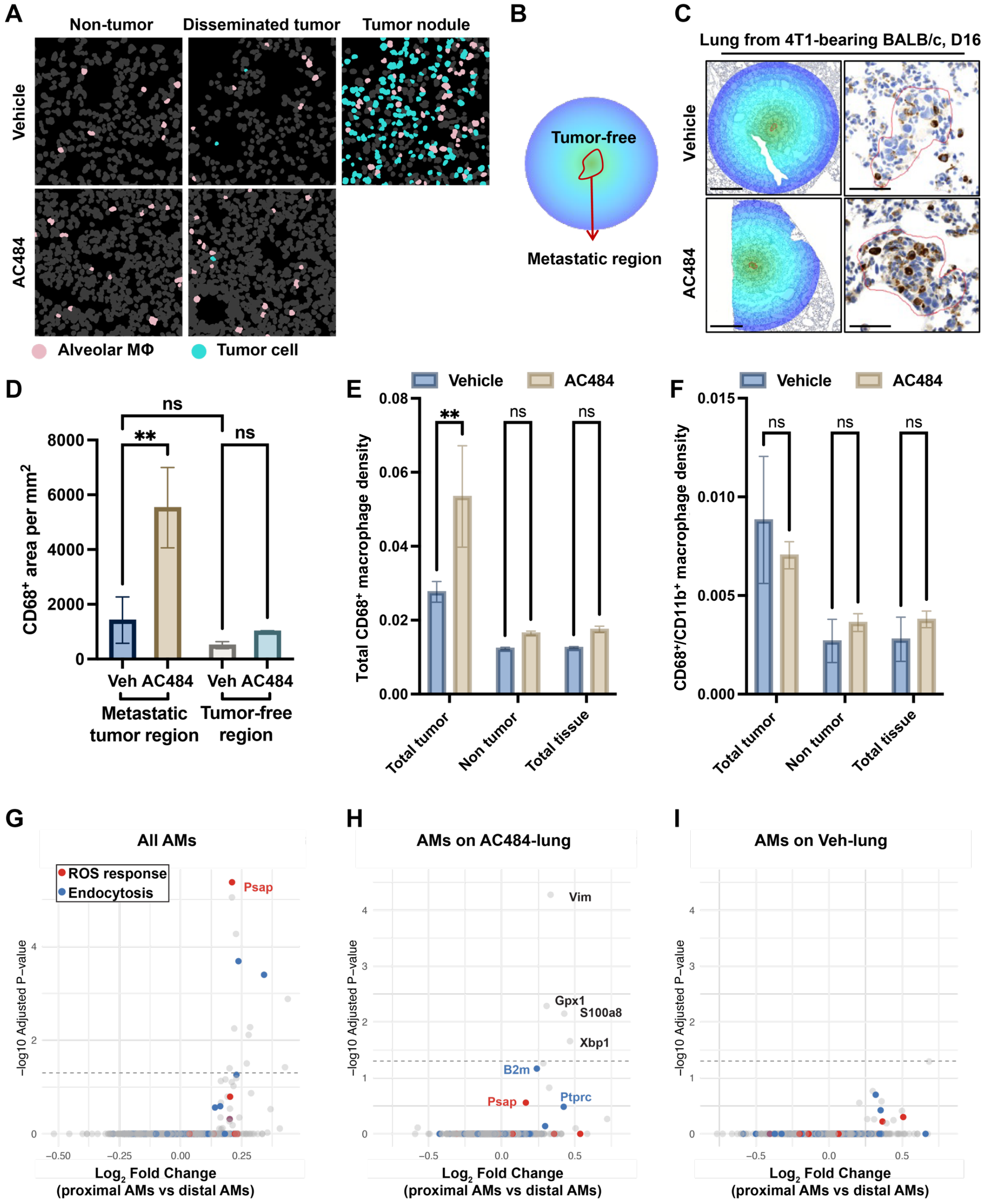

**Figure S6 (related to Figure 3). AC484 promotes macrophage infiltration at metastatic regions and affects transcription profiles in AMs**

**A**. Representative spatial distribution of AMs and disseminated tumor cells in lung sections from Figure 3C. **B.** Annotation of lung sections delineating metastatic tumor regions (red outlined) and tumor-free parenchyma (concentric blue rings) used for quantification in C. **C.** Chromogenic CD68 immunohistochemistry with hematoxylin counterstaining on lungs from 4T1-bearing mice treated from Day 7 post-implantation; tissues harvested on Day 16. Scale bar = 500 µm (left) or 50 µm (right). **D**. $CD68^+$ cell density (area/mm²) was analyzed across at least two independent metastatic regions. One-way ANOVA test. **E-F**. Quantification of total $CD68^+$ macrophages (**E**) and $CD11b^+/CD68^+$ interstitial macrophages (IMs) (**F**) across three independently immunofluorescence-stained lung sections as in Figure 3G. Unpaired t-test: ns, $p > 0.05$; * $p \leq 0.05$; ** $p \leq 0.01$; *** $p \leq 0.001$. **G-I.** Differential gene expression analysis comparing tumor-proximal and tumor-distal AMs. Tumor-proximal and -distal AMs were defined using a 300 µm cutoff, which approximates the median distance of AMs to disseminated tumor cells. The analysis was performed on all AMs across treatment groups (**G**), AMs from AC484-treated lungs (**H**), and AMs from vehicle-treated lungs (I). Genes are color-coded by the Gene Ontology (GO) pathway. Significance cutoff: BH adjusted $p < 0.05$.

#### Figure S7, Related to Figure 4

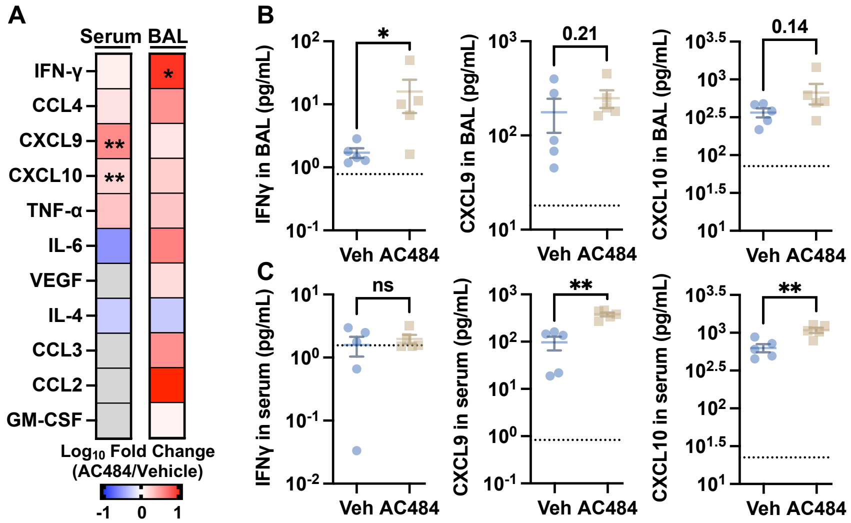


**Figure S7 (related to Figure 4). AC484 promotes IFNγ production in mouse BAL at Day 29 post 4T1 tumor inoculation**

**A.** Serum and BAL cytokine level comparison in 4T1-bearing BALB/c mice treated with AC484 or vehicle. Samples were collected on Day 29. Grey indicates values below the detection threshold. Cytokines in the 13-plex panel that did not reach the detection limit in either condition were not plotted. **B-C.** Specific cytokine levels in BAL fluid (**B**) and serum (**C**) from A. Shown are individual mice (dots) with mean ± SEM (bars). Dotted lines indicate detection limits. Unpaired t-test: ns, $p > 0.05$; * $p \leq 0.05$; ** $p \leq 0.01$; *** $p \leq 0.001$.

### Figure S8, Related to Figure 5

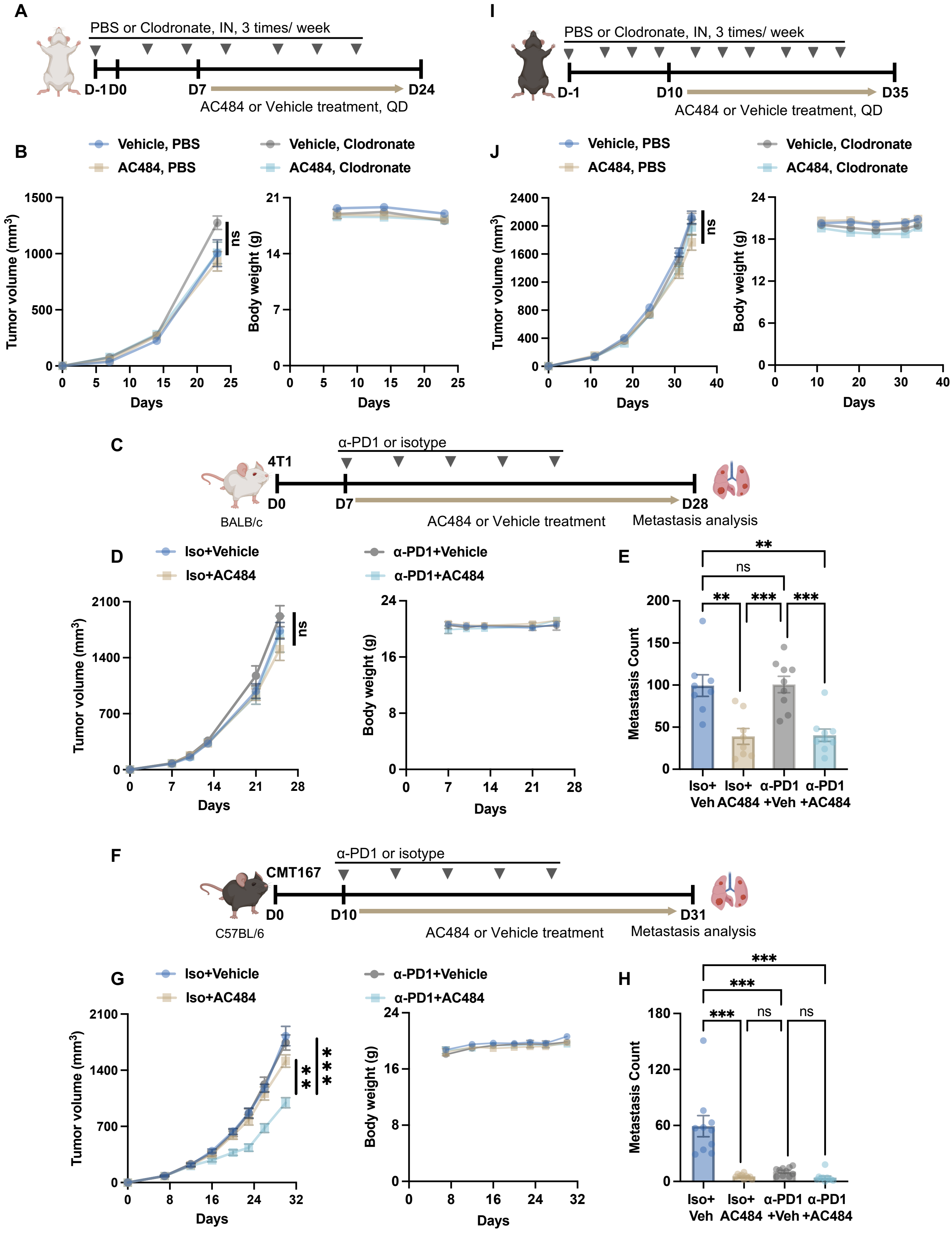

**Figure S8 (related to Figure 5). Inhibiting PTPN1/2 by AC484 suppresses metastasis in both anti-PD1-resistant and -responsive tumor models**

**A**. Experimental schedule as in Figure 5H. **B**. Primary tumor growth kinetics and corresponding body weight measurements for mice bearing 4T1 tumors in A. **C**. Experimental scheme showing BALB/c mice were implanted with 4T1 tumor cells and treated with AC484, anti-PD1 alone, their combination, or vehicle/isotype controls. **D**. Primary tumor growth and body weight from mice in C. **E**. Lung surface metastases were counted at the endpoint from mice treated in C. **F**. Experimental scheme showing C57BL/6 mice implanted with CMT167 tumor cells and treated as indicated. **G-H.** Primary tumor growth and body weight (**G**), and lung surface metastases quantification at endpoint (**H**) from mice in F. **I.** Experimental schedule as in Figure 5K. **J.** Primary tumor growth kinetics and corresponding body weight measurements for mice bearing CMT167 tumors in I. For all tumor growth kinetics, shown are mean ± SEM (bars). For all metastasis analyses, shown are individual mice (dots), with mean ± SEM (bars). N ≥ 8 mice per group. One-way ANOVA test: ns, $p > 0.05$; * $p \leq 0.05$; ** $p \leq 0.01$; *** $p \leq 0.001$.

Figure S9, Related to Figure 5

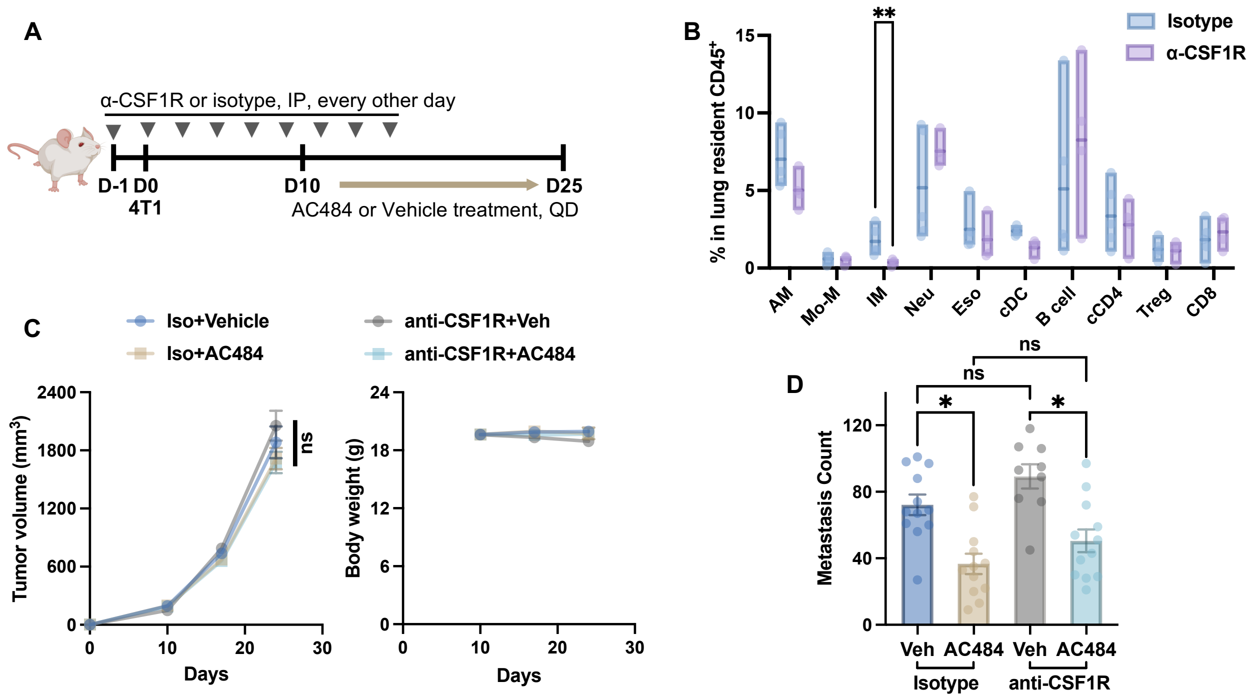


**Figure S9 (related to Figure 5). Depleting interstitial macrophage using anti-CSF1R does not affect the ability of AC484 to suppress metastasis**

**A**. Experimental schema showing BALB/c mice received 500 µg of anti-CSF1R or isotype control antibodies starting from Day -1, followed by 4T1 tumor implantation and the indicated treatments. **B**. Flow cytometry analysis of lung-resident $CD45^+$ immune cells the day after the $7^{th}$ antibody dose. **C**. Primary tumor growth and body weight of BALB/c mice treated as in A. **D**. Lung surface metastases were quantified at the endpoint in mice from A. For all bar graphs, data represent individual animals (dots) and mean ± SEM (bars). Unpaired t-test for two-group comparisons; one-way ANOVA for more than two groups: ns, $p > 0.05$; * $p \leq 0.05$; ** $p \leq 0.01$; *** $p \leq 0.001$.

#### Figure S10, Related to Figure 6

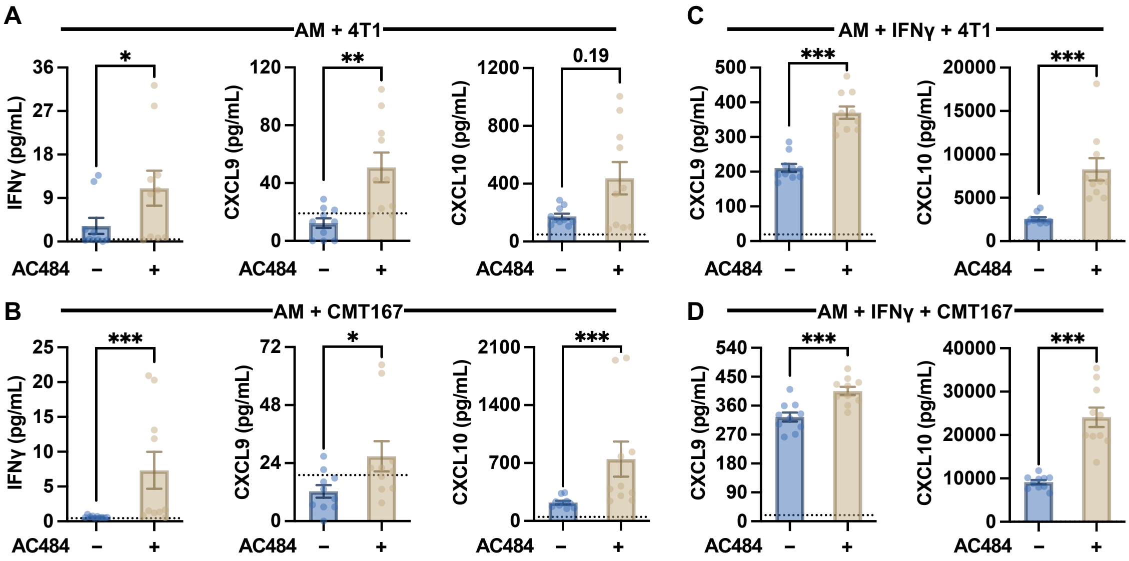


**Figure S10 (related to Figure 6). AC484 promotes IFNγ production and IFNγ-induced chemokines in AM-tumor cocultures**

**A-D.** The indicated cytokine levels in conditioned medium from AMs cocultured with 4T1 (**A, C**) or CMT167 tumor cells (**B, D**). The cocultures were performed in the absence or presence of IFNγ (1.6ng/mL) ± AC484 treatment. Shown are individual samples (dots), with mean ± SEM (bars). Dotted lines indicate detection limits. Unpaired t-test: ns, $p > 0.05$; * $p \leq 0.05$; ** $p \leq 0.01$; *** $p \leq 0.001$.

**Figure S11, Related to Figure 7**

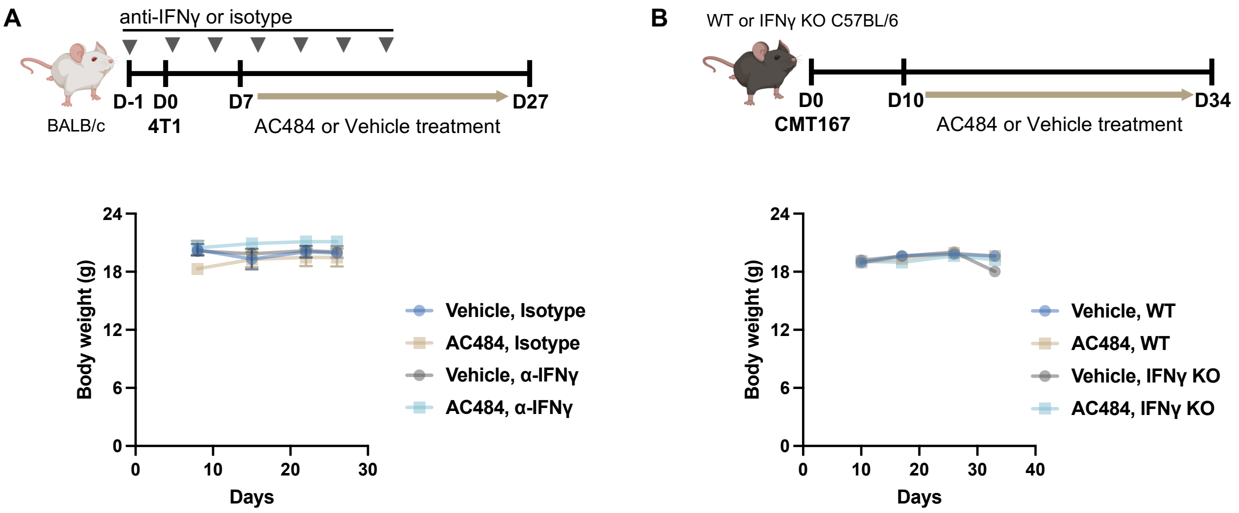


**Figure S11 (related to Figure 7). Blocking IFNγ does not affect mouse body weight with or without AC484 treatment**

**A, B.** Experimental schedules as in Figure 7A and D and body weight measurement for mice bearing 4T1 (**A**) and CMT167 (**B**) tumors.

**Supplementary Table 1: Sequences of sgRNA and primers.**

| **Name** | **Sequences** |
|---|---|
| *Ptpn1* sgRNA1 | UGAUAUAGUCAUUAUCUUCC |
| *Ptpn1* sgRNA2 | GGUGAGAAUAUAGCUCCUCU |
| *Ptpn1* sgRNA3 | CAACAGCUCUCAAGUUCUUC |
| *Ptpn2* sgRNA1 | GUCUCCCGCUCGGGCGGAAG |
| *Ptpn2* sgRNA2 | CUAAGACUCACCAAGUAUAA |
| *Ptpn2* sgRNA3 | AACCAUCGAGCGGGAGUUCG |
| Scramble negative control sgRNA | Scrambled sgRNA#1 |
| *Ptpn1* Forward and Sequencing Primer | GTCATCCTAACAGTGAGGCCC |
| *Ptpn1* Reverse Primer | TCTAGGGTCTTCTCCTGCCC |
| *Ptpn2* Forward Primer | AACACGGTCAGCGAGGAAG |
| *Ptpn2* Reverse Primer | CACAGTGCCAGCGCTCTC |
| *Ptpn2* Sequencing Primer | ATTCGGCCGACAAGAACAC |

**Supplementary Table 2: Flow cytometry antibodies**

| Specificity | Fluorochrome | Clone | Vendor |
|---|---|---|---|
| CD19 | BUV395 | 1D3 | BD Biosciences |
| CD45 | BUV661 | 30-F11 | BD Biosciences |
| CD8 | BUV805 | 53-6.7 | BD Biosciences |
| CD45 | BB700 | 30-F11 | BD Biosciences |
| FoxP3 (intracellular) | PE-Cy5.5 | FJK-16S | Thermo Fisher |
| CD103 | BV480 | 2E7 | BD Biosciences |
| CD44 | BV510 | IM7 | BioLegend |
| CD4 | BV570 | GK1.5 | BioLegend |
| CD49b | APC | DX5 | BioLegend |
| NK1.1 | APC | PK136 | BioLegend |
| CD11b | Alexa Fluor 700 | M1/70 | BioLegend |
| TCR-β | APC-Cy7 | H57-597 | BioLegend |
| Ly-6G | BUV563 | 1A8 | BD Biosciences |
| CD11b | BB515 | M1/70 | BD Biosciences |
| CD169 | PE-Dazzle 594 | 3D6.112 | BioLegend |
| XCR1 | BV421 | ZET | BioLegend |
| Ly-6C | BV570 | HK1.4 | BioLegend |
| MHC-II (I-A/I-E) | BV650 | M5/114.15.2 | BioLegend |
| CD11c | BV750 | N418 | BioLegend |
| CD172a | Alexa Fluor 700 | P84 | BioLegend |
| Zombie NIR | – | – | BioLegend |
| F4/80 | APC-Fire810 | BM8 | BioLegend |
| CD68 (intracellular) | BV711 | FA-11 | BioLegend |
| CD64 | APC | X54-5/7.1 | BioLegend |
| Siglec-F | Kiravia Blue 520 | S170007L | BioLegend |
| NKp46 | BV421 | 29A1.4 | BioLegend |

**Supplementary Table 3: Gating Hierarchy for Immune Cell Identification in Alveolar Macrophage Depletion Studies.**

| **Immune Cell Population** | **Gating Hierarchy** |
|---|---|
| Neutrophils | CD11b+/Ly-6G+ |
| Alveolar Macrophages | CD68+/CD64+/CD11b-/Siglec-F+ |
| Interstitial Macrophages | CD68+/CD64+/CD11b+/Siglec-F-/Ly-6C- |
| Monocyte-derived Macrophages | CD68+/CD64+/CD11b+/Siglec-F-/Ly-6C+ |
| Eosinophils | SSC hi/CD11b+/Siglec-F+ |
| Conventional Dendritic Cells | CD64-/CD11c+/MHC-II+ |
| B Cells | TCRβ-/CD19+/MHC-II+ |
| $CD8^+$ T Cells | TCRβ+/CD8+/CD4- |
| Conventional $CD4^+$ T Cells | TCRβ+/CD8-/CD4+/FoxP3- |
| Regulatory $CD4^+$ T Cells | TCRβ+/CD8-/CD4+/FoxP3+ |
| Natural Killer Cells | TCRβ-/NKp46+ (BALB/c) or TCRβ-/NK1.1+ (C57BL/6) |
